# Clean Breaks: Attention Force Field Deconvolution of Recombination Across Sequence Manifolds

**DOI:** 10.64898/2026.10.19.760237

**Authors:** Sergei L. Kosakovsky Pond, Danielle Callan, Steven Weaver, Hannah Verdonk, Jordan Zehr, Anton Nekrutenko, Taylor K. Paisie, Darren P. Martin

## Abstract

Homologous recombination and horizontal gene transfer assemble mosaic chromosomes across divergent lineages. Reticulate evolution violates bifurcating trees, yet existing detectors enforce a stark trade-off: phylogenetic changepoint models optimize partitioned topologies at exponential computational cost, whereas sequence triplet heuristics evaluate rapidly but suffer from severe combinatorial event overcounting. We introduce `RhizAeon`, an open-source platform that deconvolves reticulate evolution through continuous trajectory mechanics across metric embeddings in linear time (*O*(*NKL*)). Projecting aligned sequences into canonical metric coordinates and routing lineage-specific rate variation into an unpolarized root sink, the engine iteratively scrubs mosaic tracts and verifies breakpoints through Brownian-bridge statistics, dyadic matched filters, and maximum-likelihood polishing. Across canonical coalescent benchmarks and a continuous parameter sweep of 1,000,000 whole genomes, `RhizAeon` bounds false-positive rates well below the 5.0% nominal ceiling (0.58% on 1,200 rate-variation nulls and 1.01% across 333,334 continuous sweep nulls). Leave-one-in verification and bidirectional overlap clustering collapse co-recombinant clades into single ancestral reticulations. Empirical analyses across wildlife sarbecovirus Spike chimeras, multidrug-resistant *Streptococcus pneumoniae* chromosomes (2.22 Mb), and 379 primate *HLA* Class I alleles reconstruct complex whole-genome mosaicism, cassette exchanges, and micro-conversions in seconds. The core engine compiles to WebAssembly for interactive execution within standard web browsers.

## 1 Introduction

Homologous recombination, which we will hereafter refer to simply as recombination, substantially contributes to the diversification of most known viruses and cellular lifeforms. Why recombination is so rampant in the biosphere remains unsettled, though three mechanisms likely dominate. It serves as a core process in repairing double-stranded DNA breaks in cellular lifeforms, enables the “rewinding” of maladaptive evolution by purging genomes of harmful mutations, and enables the “fast-forwarding” of evolution by concentrating distinct adaptive mutations within individual genomes (Didelot and Maiden, 2010; Simon-Loriere and Holmes, 2011). The fact that biological processes have evolved to promote homologous recombination across a broad array of RNA viruses—where it plays no role in repairing broken genome strands—suggests that its primary adaptive value often lies in shuffling mutations from one generation to the next. Nevertheless recombination rates vary widely between different viruses (Simon-Loriere and Holmes, 2011) and cellular lifeforms (Didelot and Maiden, 2010; McVean et al., 2004) and, when it does occur, it tends to occur more frequently in some genome regions than in others (Lytras et al., 2022; McVean et al., 2004). Further, contingent on the precise sites in a genome where it occurs, it has a higher or lower tendency to yield recombinant offspring genomes that are either fitter or less fit than their parental genomes (Lytras et al., 2022; Simon-Loriere and Holmes, 2011).

In the context of pathogenic microbe evolution, these recombinant offspring genomes could have altered transmission dynamics, cellular tropisms, host ranges, pathogenicity, antigenicity, immune-evasiveness and antimicrobial resistance (Alfonsi et al., 2024; Croucher et al., 2011; Didelot and Maiden, 2010). In sexually reproducing eukaryotes, across successive generations highly orchestrated inter-chromosomal recombination during meiosis enables the purging of deleterious mutational load and the assembly of hybrid genomes with complex novel phenotypes (McVean et al., 2004; Rieseberg et al., 2003).

While interesting in its own right, recombination is also a nuisance in the context of phylogenetics-based molecular evolution analyses. Specifically, it undermines the core assumption of phylogenetics: that the evolutionary relatedness of a set of homologous sequences can be accurately described using a bifurcating tree. If recombinant genomes have tracts of sequence inherited from parental genomes that had sufficiently distinct evolutionary histories, attempts to place the recombinant within a single bifurcating phylogeny will tend to shorten deep inner branches, lengthen outer branches and cause spurious branching patterns (Kosakovsky Pond et al., 2006; Schierup and Hein, 2000). These recombination-induced tree distortions will at least mildly impact the downstream inference of molecular clock rates, the estimation of node dates, ancestral sequence and ancestral character state inference, transmission cluster reconstructions, and the inference of variations in historical population sizes (Posada, 2002; Schierup and Hein, 2000). When using phylogenetic codon models to analyse patterns of selection based on rates of synonymous and nonsynonymous substitutions, the apparent homoplasies caused by recombination can mimic convergent amino acid substitutions across multiple branches of a phylogenetic tree. Under standard selection detection tests, including MEME and BUSTED (Murrell et al., 2012, 2015), recombination produces spurious signals of diversifying positive selection (Kosakovsky Pond et al., 2006).

Therefore, detecting recombination breakpoint sites and identifying recombinant sequences within nucleotide sequence alignments has become a routine task in phylogenetics-based molecular evolution analyses. Identification of phylogenetically disruptive recombination breakpoint sites at the scale of whole alignments with recombination detection methods such as GARD (Kosakovsky Pond et al., 2006) can be used to partition alignments into recombination-depleted segments that can then be used individually for downstream phylogenetics-based evolutionary inference. If phylogenetically disruptive recombination breakpoints are so pervasive that recombination-depleted segments become too short for accurate phylogenetic inference, “triplet scanning” detection methods like 3SEQ (Lam et al., 2018) or those implemented in RDP5 (Martin et al., 2021) can scan every combination of three sequences in an alignment and localize breakpoints within individual sequences. With both identified recombinant sequences and breakpoint locations in hand, individual sequences can then be segmented or removed to yield full-length recombination-depleted alignments for downstream analyses.

These two different approaches to accounting for and managing the confounding effects of recombination on molecular evolutionary analyses foreshadow a much deeper divide between the ways in which the different detection methods work. GARD is an example of a phylogenetic change-point method that formulates recombination breakpoint detection as a model selection task across partitioned trees under continuous-time Markov substitution models. GARD finds optimal breakpoint boundaries through likelihood ratio criteria while fitting independent branch lengths across candidate alignment segments. Because the joint combinatorial space of partition boundaries and tree topologies scales exponentially with breakpoint count and cubically with taxon count (*O*(*L*^*K*^ · *N* ^3^) for *K* breakpoints across *N* taxa of alignment length *L*), multi-breakpoint searches across hundreds of genomes with GARD require days of cluster compute.

Triplet scanning detection methods like those implemented in 3SEQ and RDP5, forgo computationally expensive evolutionary modeling. Instead they rely on the heuristic detection of localized relative or absolute differences in the similarities of sequence pairs within the context of individual sequence triplets drawn from a bigger alignment. While evaluating rapidly across alignments of a few hundred sequences, exhaustive triplet scanning approaches scale poorly *O*(*N* ^3^*L*) to the point that they become untenable when alignment sizes increase to over a few thousand sequences. Because individual sequence triplets displaying evidence of recombination have no phylogenetic reference frame against which to decide which sequence is the recombinant (and which are parent-like), triplet scanners must be augmented with downstream phylogenetic methods to make these calls. Accurate parent-recombinant calls become particularly difficult when close relatives of both parental lineages are absent from an analyzed alignment. Downstream phylogenetic methods and other techniques (such as iteratively deconstructing or masking detected recombinant sequences and re-scanning for recombination) are also needed to determine the counts of actual ancestral recombination events underlying detected signals. Without this step, any given ancestral recombination event may be independently detected across tens or hundreds of different sequence triplets drawn from an alignment. Finally, if triplet scanning methods do not take steps to phylogenetically contextualize detected recombination signals they run the risk of identifying lineage-specific increases in substitution rates within defined genome sub-regions as evidence of recombination. All of this additional phylogenetic contextualization incurs extra computational costs over-and-above the triplet-scanning recombination signal detection components of 3SEQ and RDP5.

Here we introduce `RhizAeon`, an extremely fast and linearly scaling open-source recombination detection tool that retains phylogenetic context without the construction of discrete phylogenetic trees. Its name derives from the Greek *rhiza* (root) and evokes the interconnected network of the biological rhizome across deep evolutionary time (*aeon*) to supplant strictly bifurcating arboreal trees.

## 2 The RhizAeon Framework: Principles of Operation

`RhizAeon` models recombination through continuous trajectory mechanics and zero-drift force field dynamics in linear time (*O*(*NKL*) for *N* taxa, an alignment length *L*, and a set of *K* dynamically determined reference sequences that we refer to here as “landmark” lineages). By measuring instantaneous sequence affinity against reference lineages and absorbing lineage-specific rate variation into an unpolarized ancestral root sink, `RhizAeon` transforms recombination detection into a physical trajectory deconvolution problem. Clonal evolution produces balanced “thermal” fluctuations that cancel out as a centered Brownian bridge. Recombination breaks this physical equilibrium and imparts sustained directional momentum that pinpoints breakpoint sites while consolidating sets of recombination signals attributable to the “co-recombinant” descendants of ancestral recombinant sequences into single ancestral recombination events.

Continuous trajectory tracking across sequence manifolds differentiates recombination signals from signals of clonal divergence without a discrete tree search. Scanning along an alignment from 5’ to 3’, a non-recombinant genome maintains a stable genetic identity: its closest relatives remain constant and trace parallel paths through sequence space. Recombination disturbs this stability. A mosaic genome merges genetic material from two or more distinct parental lineages. In principle, a single recombination crossover lacks intrinsic biological polarity. If a recombinant sequence combines a segment from parental lineage *A* with a segment from parental lineage *B*, sites within the *A*-derived segment view *A* as the recipient background and *B* as the foreign introgression, whereas sites within the *B*-derived segment view the roles in reverse. For consistency across the framework, we define the *home* lineage (or major parent) of a focal sequence as the reference lineage exhibiting the highest cumulative affinity across the full alignment length, and the *donor* lineage (or minor parent) as the reference lineage toward which affinity shifts within a localized recombinant tract. As a scan progresses along a mosaic genome, the sequence travels alongside its home lineage, then abruptly shifts affinity toward a donor lineage across the recombinationally acquired tract. To capture these shifts in affinity without constructing trees or using sliding windows, `RhizAeon` measures the instantaneous affinity of each query sequence against a representative panel of landmark reference sequences at every polymorphic site of an alignment.

To evaluate affinity at each polymorphic site, `RhizAeon` adapts cross-attention from transformer neural networks. Each focal genome serves as the query, while the landmark reference sequences act as keys. At every position along the alignment, the query evaluates which landmark reference sequence it resembles to assign local evolutionary affinity. Rather than treating all nucleotide matches equally, continuous-time Markov substitution probabilities weight each attention score: sharing a rare derived allele commands strong attention, whereas common transitions command less. To prevent private mutations from creating spurious matches, the landmark panel includes an unpolarized ancestral root token that absorbs alleles absent from every contemporary reference. A softmax operator normalizes these scores into a probability distribution that sums to one across all landmark reference sequences and the root token at each site. Because individual polymorphic sites exhibit stochastic site-to-site variation in nucleotide substitution patterns and instantaneous attention weights, `RhizAeon` avoids evaluating positions in isolation or smoothing them with fixed sliding windows. Instead, it subtracts each landmark reference sequence’s alignment-wide mean attention to integrate cumulative deviation along the alignment. In non-recombinant regions, positive and negative site fluctuations cancel out and wander near zero like a random walk. When a scan along a sequence enters a recombinant tract, sustained attention toward the donor reference drives the cumulative trajectory steadily upward until donor attention ceases at the breakpoint junction and the curve descends. A single breakpoint forms a sharp triangular arch, whereas an inserted cassette produces a trapezoidal plateau. The switch points fall at the peaks and inflection points of these cumulative attention curves to localize breakpoints to exact nucleotide coordinates. A physical force field governs the transition from site-by-site attention to continuous trajectories.

At each polymorphic site, the attention directed toward a landmark reference sequence acts as an attractive force that pulls the query sequence toward that reference in metric space. Subtracting the sequence’s alignment-wide baseline attention establishes a zero-drift force field in which positive and negative pulls average to zero across the sequence. Under clonal evolution, mutations arise independently at random positions. Instantaneous forces fluctuate in opposing directions and cancel out like thermal Brownian motion, confining the sequence trajectory within its home corridor. Recombination breaks this physical equilibrium. Within an introgressed segment, every shared derived allele exerts a sustained directional pull toward the donor lineage. Sustained donor attraction imparts momentum to the sequence trajectory and drives a macroscopic excursion away from the home corridor across sequence space. The physical breakpoints correspond to the exact genomic coordinates where the external directional force turns on and off.

Different modes of recombination generate distinct geometric signatures within the force field. In a single breakpoint recombination event, a sequence begins in its home corridor and, at the breakpoint site, switches into a donor corridor for the remainder of the recombinant sequence. The directional force acts persistently from the breakpoint onward and drives cumulative momentum to a solitary peak at the breakpoint junction before returning to zero at the end of the recombinant sequence. The resulting profile forms a triangular arch whose vertex pinpoints the breakpoint location. For a double breakpoint recombinant (hereafter referred to as a cassette), the directional force turns on at the 5’ breakpoint and turns off at the 3’ breakpoint to form a trapezoidal plateau: a steep upward climb at the first breakpoint, a sustained plateau across the cassette, and a steep descent at the second breakpoint.

When the donor lineage that provided the tract of sequence acquired by a recombinant remains unsampled, the introgressed segment exhibits a sharp loss of affinity to the home lineage without pointing toward any sampled contemporary taxon in the alignment. Rather than manufacturing a false pairing with an arbitrary contemporary leaf or silently discarding the signal, `RhizAeon` detects the localized departure from the home lineage and explicitly flags and labels the introgressed segment as originating from an unsampled `“Ghost”` donor lineage. This enables investigators either to retain the event as a basal/unsampled introgression or to mask and strip the ghost-derived tract prior to downstream phylogenetic inference, while achieving zero false framing of contemporary background parents. When a close relative of the unsampled donor is present in the dataset (a surrogate sister lineage), trajectory tracking captures the shared derived alleles inherited from their common ancestor and attributes the transfer to the sampled surrogate sister reference (achieving 88.3%–100.0% surrogate donor attribution accuracy across divergence cohorts; Section 3.2).

Localizing breakpoints combines the multi-scale speed of the force field with the fine-scale precision of phylogenetic likelihood. In natural genomes, recombination tract lengths vary by orders of magnitude. Fixed sliding windows can sometimes fail across this span: narrow windows might succumb to local mutational noise, whereas wide windows might blur signals generated by short recombination tracts. Continuous dyadic wavelet filters eliminate this trade-off by scanning cumulative momentum across a geometric ladder of scales from fifteen to five hundred base pairs. Fine-scale wavelets isolate short recombination tracts, while broad-scale wavelets lock onto larger exchanges without distortion from localized rate fluctuations. The zero-crossings and extrema of these wavelet responses narrow candidate breakpoint sites from whole alignments down to a localized window of fifteen to thirty base pairs.

In the second stage, profile likelihood polishing refines each candidate window to individual nucleotide coordinates. Because mutations are unevenly distributed across genomes, the identified breakpoint site can lie anywhere between adjacent informative mutation sites. Within the candidate window, a trial boundary slides nucleotide by nucleotide across the three-sequence triplet comprising the candidate recombinant sequence, its resolved home reference sequence, and its resolved donor reference sequence. Sites on the home side of the trial boundary are evaluated under the pairwise substitution likelihood to the home reference, while sites on the donor side are evaluated under the substitution likelihood to the donor reference. The optimal breakpoint is placed at the exact coordinate that maximizes this joint triplet profile likelihood. Furthermore, the curvature of the likelihood surface yields rigorous confidence intervals: when parental lineages share an identical conserved stretch, the polisher reports the full uninformative plateau rather than manufacturing an artificial single-base prediction.

In cohorts where recombination is pervasive, an initial static selection of reference landmarks risks incorporating mosaic sequences into the reference coordinate frame, which can obscure secondary recombination tracts or invert parent and recombinant identities. To prevent reference contamination without discarding sequences, `RhizAeon` employs a two-pass iterative forward scrubbing and backward leave-one-in verification architecture. During the forward pass, candidate recombinant segments are ranked from highest to lowest signal-to-noise ratio and iteratively scrubbed by replacing the detected recombinant tract with gap characters in the working alignment. Outside its gap-masked tract, every sequence remains intact and continues to serve as an uncorrupted landmark reference for the rest of the cohort. Within the cross-attention force field, gap-masked query sites exert zero net force (Δ*F* = 0), preventing artificial drawdown decay across scrubbed intervals while preserving natural alignment indel distances across unmasked sites. Once the forward pass converges to a recombination-depleted landmark alignment free of trajectory perturbations, a backward leave-one-in verification pass restores each candidate recombinant segment one at a time in isolation while keeping all other recombinant tracts masked. Each restored segment is re-evaluated for parental affinities and re-polished across its boundaries without interference from overlapping or adjacent breakpoint signals. Restoring a segment can also reveal a reciprocal parent-recombinant polarity inversion, an algorithmic symmetry artifact where donor *D* initially tested positive as a putative recombinant between *R* and *H*. Whenever treating *R* as the recombinant yields a sharper transition contrast and lower flanking strain, the engine swaps the assignment, scrubs the true recombinant tract in *R*, and leaves parental sequence *D* intact.

Before certifying any candidate recombination event in the backward verification pass, an orthogonal three-stage audit evaluates the raw alignment sites. First, a Poisson informative-site floor requires every candidate tract to contain at least three unambiguous informative sites where the donor allele differs from the home allele and matches the recombinant. Second, when a candidate tract contains a high density of mutations separating the query sequence from its home reference that are absent from all other sampled sequences, the engine tests their spatial distribution. Concentrated divergence blocks from a basal unsampled outgroup are retained and labeled as unsampled `“Ghost”` donor introgressions, whereas dispersed private substitutions are absorbed into the root sink and rejected. Third, for tracts attributed to a sampled donor reference, an orthogonal Fisher’s exact test evaluates a two-by-two contingency table across the candidate recombinant, resolved home reference, and resolved donor reference triplet. This test compares the counts of polymorphic sites where the recombinant matches the donor versus the home reference inside the candidate tract against those in the flanking alignment regions (excluding monomorphic sites and positions where the home and donor references carry identical nucleotides), and must achieve significance under family-wise error rate control across the cohort. Together, iterative scrubbing, leave-one-in verification, and raw-site auditing maintain strict false-positive control even under extreme mutation rate heterogeneity.

While many detected recombinants in sparse empirical surveys appear as singletons, expanding viral outbreaks and bacterial clones frequently produce clusters of multiple sampled descendants that inherited the same mosaic architecture from a common recombinant ancestor. When multiple descendants of an ancestral recombinant are sampled within an alignment, graph adjudication clusters these co-recombinant sequences whenever they share concordant parental lineages and overlapping breakpoint coordinates, consolidating the group into a single ancestral recombination event. Defining one parent as “home” (major) and the other as “donor” (minor) becomes arbitrary whenever crossover tracts span a large fraction of the alignment. Ancestral consolidation therefore evaluates both direct (Home_*A*_ ≈ Home_*B*_, Donor_*A*_ ≈ Donor_*B*_ with overlapping tracts) and inverted (Home_*A*_ ≈ Donor_*B*_, Donor_*A*_ ≈ Home_*B*_ with complementary breakpoint junctions) parental polarities so that co-recombinant isolates with swapped major/minor parent labels are grouped into the same ancestral event.

## 3 Coalescent Simulation Benchmark and Operational Evaluation

### 3.1 Coalescent ARG Benchmark and Calibration Against 3SEQ

To establish a baseline against 3SEQ—evaluated here as a fast, widely used representative among exact triplet-scanning algorithms—we benchmarked `RhizAeon` across the 3,200 coalescent simulation scenarios of Posada and Crandall (2001) (*N* = 10 taxa, *L* = 1,000 nt, Figure 1, Supplementary Table S2). Across 1,200 non-recombinant alignments simulated with site-to-site mutation rate variation (*α* ∈ {2.0, 0.5, 0.05}), both algorithms maintained strict false positive control well below the nominal 5% ceiling (Figure 1C). On these small 10-taxon recombination-free alignments, 3SEQ recorded a 0.17% false positive rate (2 / 1,200 trials, 95% Wilson score interval [0.05%, 0.60%]). `RhizAeon` yielded 0.58% (7 / 1,200 trials, 95% Wilson score interval [0.28%, 1.20%]), with 0.0% false positives across all 600 trials at *θ* ∈ {10, 50} and across 8 of the 12 rate-variation conditions. Occasional false alarms occurred primarily at high sequence divergence (*θ* = 200, accounting for 6 of the 7 false positives across *α* ∈ {2.0, 0.5, 0.05}) due to stochastic homoplasy across the 1,000 nt locus, instead of concentrating under extreme site-to-site rate variation. Because the Posada and Crandall (2001) protocol models Gamma rate variation strictly across sites rather than heterotachously along individual lineages, 3SEQ remains well-calibrated in this regime. When lineage-specific rate accelerations are introduced (Section 3.3), private mutations on a single branch lack directional affinity toward candidate donors and route harmlessly into `RhizAeon`’s root sink.

**Figure 1.**
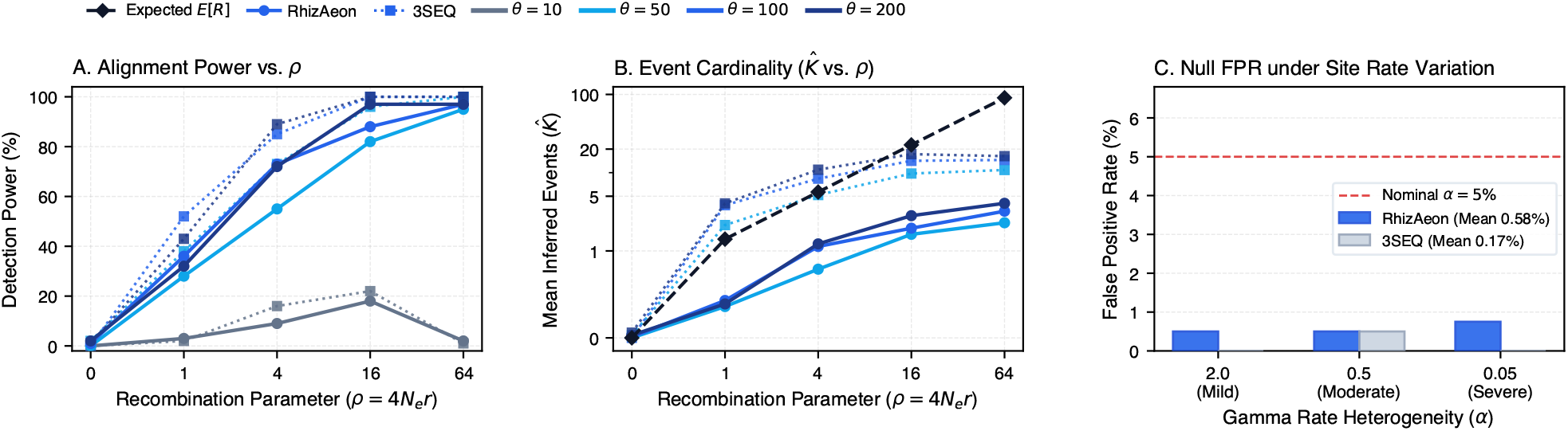
Performance of `RhizAeon` and 3SEQ across the 3,200-trial Posada and Crandall (2001) coalescent benchmark (*N* = 10, *L* = 1,000 nt). (A) Alignment-level detection power across population recombination rates (*ρ* = 4*N*_*e*_*r* ∈ [0, 64]) and mutation densities (*θ* = 4*N*_*e*_*µ* ∈ [10, 200]). (B) Inferred event cardinality 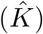 versus Hudson’s theoretical expectation (E[*R*], black dashed line), contrasting unconsolidated 3SEQ triplet counts against consolidated `RhizAeon` events. (C) Empirical false-positive rate (FPR, %) across 1,200 non-recombinant alignments under site-to-site Gamma rate variation (*α* ∈ {2.0, 0.5, 0.05}; red dashed line marks the 5% nominal threshold).

Across the 1,600 recombinant alignments (*ρ* ∈ {1, 4, 16, 64}), detection power depended primarily on mutational availability (Figure 1A). In low-diversity datasets (*θ* = 10, fewer than 25 variable sites), sensitivity was strictly limited by mutational density: both `RhizAeon` and 3SEQ detected 1.0%–22.0% of alignments across *ρ* ∈ [1, 64], where frequent recombination events split sequences into short segments carrying few mutations. At higher mutational densities (*θ* ≥ 50), 3SEQ’s more permissive default triplet threshold provided a sensitivity advantage at intermediate recombination rates (e.g., 73.0% vs. 55.0% at *θ* = 50, *ρ* = 4), while both methods converged to 88.0%–100.0% power at *ρ* ≥ 16 (*θ* ∈ {100, 200}). Even on compact 10-taxon alignments where 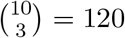 triplets require minimal combinatorial work, `RhizAeon` executed in a mean of 13.2–26.2 ms per alignment across *θ* ∈ [10, 200] (17.4 ms across all 1,200 rate-variation nulls), compared to 27.6–36.8 ms for 3SEQ. As cohort sizes expand to *N* = 32–55 taxa in the surveillance benchmarks below, `RhizAeon`’s O(*NKL*) trajectory architecture runs up to 5.5-fold faster than *O*(*N* ^3^*L*) triplet enumeration.

A central distinction between `RhizAeon` and raw 3SEQ outputs lies in event cardinality and bipartite event precision (Figure 1B and Supplementary Table S2). Because our Hudson coalescent simulator explicitly records the simulated Ancestral Recombination Graph (ARG) and extracts all observable topological transfers (*K*_true_, Section S7), we directly compared inferred event counts 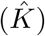 against both the exact simulated ARG events and Hudson’s theoretical expectation (E[*R*], dashed line). When 3SEQ is run without downstream post-processing, it outputs every sequence triplet displaying a significant hypergeometric random-walk excursion. Even when an alignment contains only a single recombinant sequence, multiple combinations of parent-like proxy sequences in the alignment—compounded by co-recombinant descendants inheriting the same ancestral crossover—cause raw triplet counts to reach (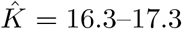 at *θ* = 200, *ρ* ∈ {16, 64}. In contrast, `RhizAeon` analyzes all sequences jointly through iterative scrubbing and leave-one-in verification, grouping shared segments into consolidated ancestral events ( 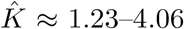 at *θ* = 200, *ρ* ≥ 4). Under strict one-to-one Hungarian bipartite matching weighted by parental triad attribution (*S*_triad_) and SNP overlap (*J*_SNP_, Equation 10), `RhizAeon` achieves 2.3-to 2.8-fold higher mean continuous event-level precision than unconsolidated 3SEQ across *θ* ≥ 50 (36.5%–63.2% vs. 4.6%–46.5% at *θ* = 200). However, as emphasized in the 3SEQ documentation (Lam et al., 2018), raw 3SEQ triplet lists are intended to be post-processed via downstream consolidation scripts that deduplicate redundant proxy-parent triplets and group overlapping breakpoint intervals. We therefore implemented a faithful downstream 3SEQ consolidation pipeline—`3SEQ (Consolidated)`—and evaluated it head-to-head against both raw `3SEQ` and `RhizAeon` across four specialized epidemiological and evolutionary benchmark suites.

### 3.2 Performance Across Specialized Surveillance and Evolutionary Regimes

Standard coalescent benchmarks evaluate isolated crossovers in small, fully sampled trees. In natural populations and disease outbreaks, however, recombination violates these idealized conditions (Figure 2). Recombinant lineages expand into co-recombinant transmission clusters, acquire sequence tracts from multiple distinct donors, inherit material from unsampled reservoirs, or exchange short micro-conversion tracts. To evaluate breakpoint localization, parent attribution, and cluster consolidation fairly, we compared `RhizAeon` against both unconsolidated `3SEQ` and post-processed `3SEQ (Consolidated)` across 2,280 alignments spanning four specialized regimes.

**Figure 2.**
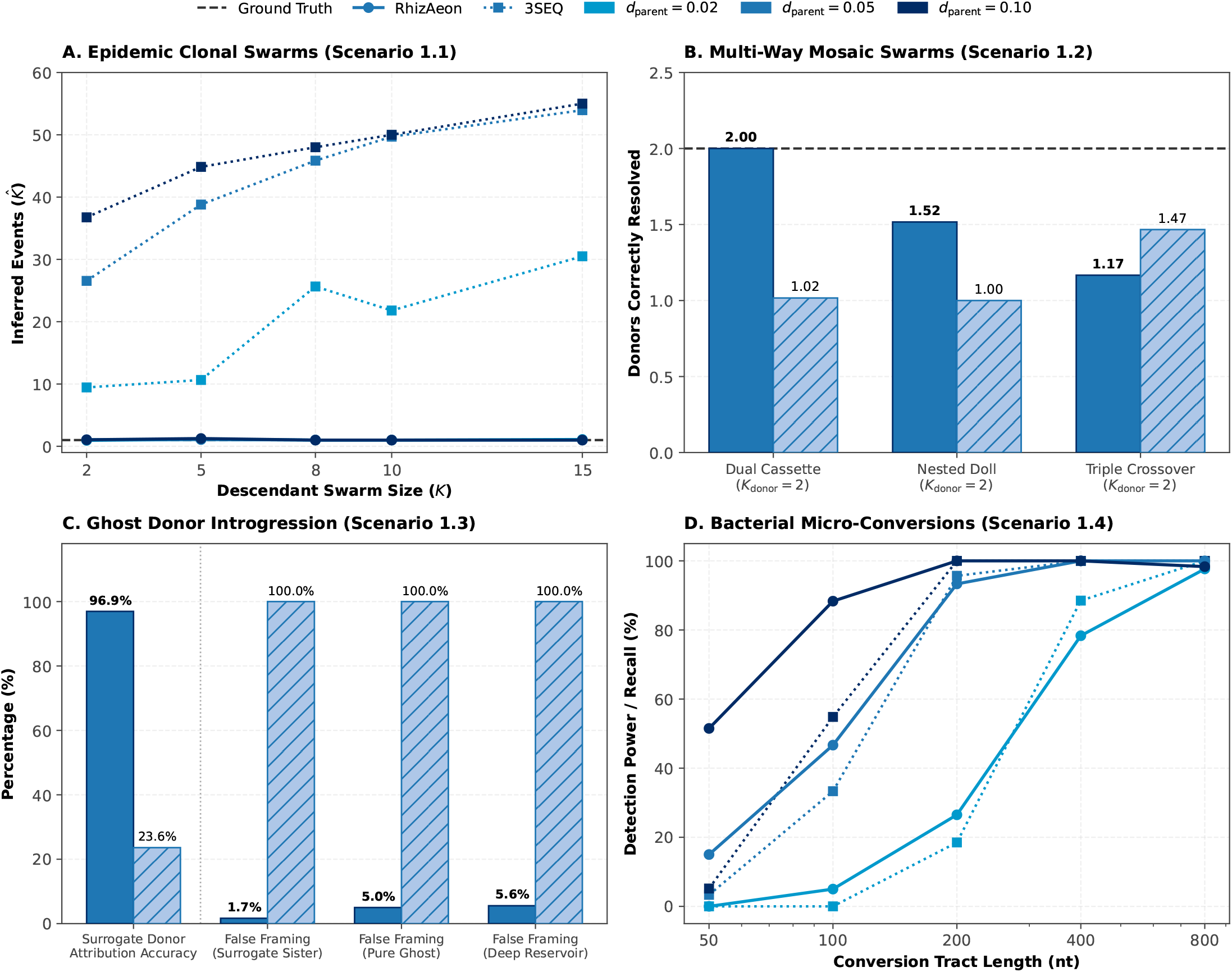
Comparison of `RhizAeon`, unconsolidated `3SEQ (Raw)`, and post-processed `3SEQ (Consolidated)` across four specialized genomic and epidemiological regimes. Columns display the simulated topology (Col. 1), primary diagnostic metric (Col. 2), breakpoint boundary overlap or error (Col. 3), and inferred event cardinality 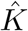 across co-recombinant cluster sizes *K* (Col. 4). Epidemic clonal expansion of a single ancestral 1.5 kb cassette into *K* ∈ {2–15} sampled descendants. (B) Multi-way mosaic clusters combining two distinct donors (*K*_donor_ = 2.0) across Dual Cassette, Nested Doll, and Triple Crossover architectures. (C) Ghost donor introgression under Surrogate Sister, Pure Ghost, and Deep Reservoir sampling regimes. (D) Bacterial micro-conversions across replacement tract lengths *L*_tract_ ∈ [50, 800] nt.

#### Epidemic Clonal Expansion

During viral outbreaks and bacterial epidemics, an ancestral recombinant frequently transmits across multiple hosts and expands into a clade or cluster of *K* sampled co-recombinant isolates (Figure 2A). Across 300 simulated outbreak cohorts (*N* = 42–55 taxa, *L* = 5,000 nt, *K* ∈ {2, 5, 8, 10, 15}), unconsolidated `3SEQ` reported every significant sequence triplet independently. Raw triplet counts inflated from 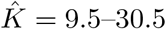 (*d*_parent_ = 0.02) to 36.8–55.0 (*d*_parent_ = 0.10), while reciprocal triplet inversions flagged background parental lineages as recombinants. Downstream consolidation in `3SEQ (Consolidated)` filters reciprocal parent inversions via directional transition contrast and clusters overlapping co-recombinant intervals. At moderate-to-high divergence (*d*_parent_ ≥0.05), this post-processing resolved the single-event epidemic regime (Figure 2A). It collapsed the co-recombinant cluster into 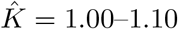 unique events with 99.0%–100.0% recall, 93.8%–100.0% precision, ≤ 2.5% background false alarms, and 5.6–14.0 nt mean breakpoint error (*J* = 0.98–0.99). `RhizAeon` achieved comparable consolidation and accuracy directly within its native linear-time pass (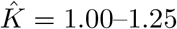 at *d*_parent_ ≥ 0.05 and 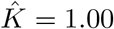 across all *K* ∈ {8, 10, 15} at *d*_parent_ ∈ {0.02, 0.05, 0.10}). It recovered 100.0% of co-recombinant descendants across all 10 conditions at *d*_parent_ ≥ 0.05 (95.0%–100.0% at *d*_parent_ = 0.02, with 100.0% in 14 of 15 conditions overall) at 92.3%–100.0% precision (100.0% in 10 of 15 conditions) and 0.00% background false positives in 11 of 15 conditions. Mean breakpoint error was 12.4–39.3 nt at *d*_parent_ = 0.10 (*J* = 0.95–0.98) and 22.3–35.2 nt at *d*_parent_ = 0.05 (*J* = 0.95–0.97), with scans completing in 2.05–6.46 s versus 1.06–10.87 s for 3SEQ. At low parental divergence (*d*_parent_ = 0.02), `RhizAeon` recovered a substantially higher fraction of co-recombinant cluster members than `3SEQ (Consolidated)` as cluster size grew (100.0% recall vs. 28.0%–45.0% for *K* ∈ {8, 10, 15}).

#### Multi-Way Mosaic Clusters

Natural viral and bacterial recombinants frequently acquire genetic material from more than one parental donor, forming multi-way mosaic architectures including dual cassettes, nested “Russian doll” transfers, and serial triple crossovers (Figure 2B). `RhizAeon` accommodates multi-parent mosaicism by iteratively scrubbing detected segments and tracking directional excursions against all landmark references simultaneously. A single query genome can thus be decomposed into multiple distinct donor transitions. In Dual Cassette recombinants (*P*_2_ and *P*_3_ cassettes embedded within a *P*_1_ background), `RhizAeon` recovered both parental donors in nearly every trial, averaging 1.97 of 2.00 expected donors overall (1.90–2.00) and 2.00 across all conditions at *d*_parent_ ≥ 0.05 (Figure 2B). Across all Dual Cassette conditions, recall reached 100.0% with 97.1%–100.0% precision (100.0% in 7 of 9 conditions), ≤ 1.6% background false alarms (0.0% across all *K* at *d*_parent_ = 0.10), and high boundary overlap (*J* = 0.95–0.97 at *d*_parent_ = 0.10). In Nested Russian Dolls (*P*_1_ → *P*_2_ → *P*_3_ → *P*_2_ → *P*_1_) and Triple Crossovers (*P*_1_ → *P*_2_ → *P*_3_), `RhizAeon` resolved an average of 1.56 (1.40–1.75) and 1.04 (0.70–1.35) donors respectively while maintaining 95.0%–100.0% recall, 95.0%–100.0% precision, and ≤ 1.7% background false positives (0.0% in 13 of 18 conditions). By contrast, 3SEQ’s hypergeometric scan extracts a single dominant recombinant segment per triplet call. Even after downstream consolidation, `3SEQ (Consolidated)` recovered only one donor per mosaic genome (1.00–1.15 of 2.00 expected donors in Dual Cassettes and 1.00 in Nested Dolls, Figure 2B). Furthermore, when a mosaic genome carries tracts from two distinct donors (*P*_2_ and *P*_3_), evaluating triplet (*R, P*_1_, *P*_2_) across the *P*_3_-derived segment distorts the flanking baseline. Consequently, `3SEQ (Consolidated)` recorded 16.2%–32.9% background false-positive calls in Dual Cassettes at *d*_parent_ ≥ 0.05 (down from 47.1%–81.0% in raw 3SEQ, compared to 0.0%–0.6% for `RhizAeon`).

#### Ghost Donor Introgression

In empirical surveillance, recombination frequently involves donor lineages absent from the analyzed alignment, such as unsampled wildlife reservoirs or extinct ancestral variants (Figure 2C). In this regime, 3SEQ does not claim to identify the true unsampled parent. Instead, it uses sampled sequences in the alignment as proxy parents to detect the localized shift in sequence similarity and localize breakpoint boundaries. When a close relative of the unsampled donor is present in the cohort (the Surrogate Sister regime), both `RhizAeon` and `3SEQ (Consolidated)` perform well. At *d*_parent_ ≥ 0.05, `RhizAeon` achieved 100.0% recombinant recall, 100.0% precision, 0.0% background and parent false alarms, *J* = 0.86–0.99 boundary overlap (0.96–0.99 in 5 of 6 conditions), and 100.0% surrogate sister attribution (84.0%–99.0% recall, 90.6%–96.3% precision, and 85.0%–100.0% attribution at *d*_parent_ = 0.02). Under the same conditions, `3SEQ (Consolidated)` achieved 100.0% recall, 88.7%–100.0% precision, *J* = 0.97–0.99, and 95.0%–100.0% surrogate sister attribution, vastly improving upon unconsolidated 3SEQ (9.3%–39.3% precision and 37.5%–56.4% background false alarms). However, when the introgressing donor branches basal to the MRCA of the sampled alignment with no close sampled relative (the Pure Ghost and Deep Reservoir regimes), a critical trade-off emerges. Unrooted 3SEQ triplets lack an outgroup reference to orient which sequence diverged. An introgressed basal ghost tract in recombinant *R* (within clade *P*_1_) makes *R* appear as a distant outgroup inside the tract, while non-recombinant *P*_1_ sequences remain close to *P*_1_ inside the tract and to proxy *P*_2_ outside it. Both raw `3SEQ` and `3SEQ (Consolidated)` therefore localize the breakpoint coordinates accurately (*J* = 0.95–0.99) but invert the recombinant identity. They fail to identify *R* as the recombinant (0.0% recall for `3SEQ (Consolidated)`) and instead flag non-recombinant *P*_1_ leaves in 50.0% of background taxa (100.0% parent false alarm rate). Instead of discarding basal introgressions or misattributing them to contemporary proxy clades, `RhizAeon`’s iterative scrubbing and orphan-divergence audit flags tracts where a sequence departs sharply from its home clade without matching any sampled reference, labeling the donor as `“Ghost”`. Across Pure Ghost cohorts at *d*_parent_ ≥ 0.05 (*K* ∈ {2, 5, 10}), `RhizAeon` detects 85.0%–100.0% of ghost-introgressed recombinants (80.0%–96.5% at *d*_parent_ = 0.02) with 95.0%–100.0% precision (100.0% in 5 of 6 conditions at *d*_parent_ ≥ 0.05), *J* = 0.83–0.99 boundary overlap, 85.0%–100.0% `“Ghost”` donor attribution, and 0.0% background false alarms in 5 of 6 conditions (Figure 2C). Similarly, across Deep Reservoir cohorts (*d*_reservoir_ = 0.15), `RhizAeon` achieves 80.0%–100.0% recall (90.0%–100.0% in 8 of 9 conditions) and 80.0%–100.0% `“Ghost”` donor attribution. At *d*_parent_ ≥ 0.05, precision reaches 90.0%–100.0% (100.0% in 5 of 6 conditions; 76.8%–95.0% at *d*_parent_ = 0.02) with *J* = 0.83–0.98 boundary overlap and 0.0% background false alarms in 5 of 6 conditions.

#### Bacterial Micro-Conversions

In bacterial transformation (e.g., *Streptococcus pneumoniae, Helicobacter pylori* ), homologous replacement frequently involves short micro-conversion tracts spanning *<* 10% of the locus (Figure 2D). Detecting micro-conversions is governed by the information-theoretic count of distinguishing polymorphisms within the tract. At *L*_tract_ = 50 nt (*L* = 5,000 nt), simulated tracts carried a mean of only 0.8–1.4 (*d*_parent_ = 0.02), 1.8–2.9 (*d*_parent_ = 0.05), and 4.9–5.2 (*d*_parent_ = 0.10) mutations. Both `RhizAeon`’s windowless dyadic matched filter and 3SEQ’s exact hypergeometric scan enforce strict family-wise error rate thresholds across all sequence comparisons. When 50 nt tracts carried fewer than two mutations (*d*_parent_ = 0.02), no method called recombination (0.0% recall and 0.0% false positives). As mutational density rose to 2.8–5.2 SNPs at *L*_tract_ = 50 nt (*d*_parent_ ∈ {0.05, 0.10}), `RhizAeon`’s multi-scale matched filter detected 15.0%–30.0% (*d*_parent_ = 0.05, *K* ∈ {2, 5}) and 50.0%–54.5% (*d*_parent_ = 0.10) of micro-conversions with 100.0% precision and 0.0% background false alarms, compared to 0.0%–10.5% for raw `3SEQ` and 0.0%–5.0% for `3SEQ (Consolidated)`. At *L*_tract_ = 100 nt (*d*_parent_ = 0.10, 8.0–10.6 SNPs), `RhizAeon` reached 80.0%–100.0% recall (mean 88.3%) with 100.0% precision and 0.0% background false alarms, exceeding both raw `3SEQ` (45.0%–70.5%) and `3SEQ (Consolidated)` (20.0%–45.0%). Across all 45 Micro-Conversion conditions (*L*_tract_ ∈ [50, 800] nt), `RhizAeon` maintained 100.0% precision and 0.00 % background false alarms. For tract lengths *L*_tract_ ∈ {200, 400, 800} nt (*d*_parent_ ≥ 0.05, ≥ 9.3 SNPs), `RhizAeon` achieved 85.0%–100.0% recall (100.0% in 15 of 18 conditions) with 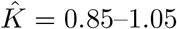 inferred events (1.00 in 14 of 18 conditions). Mean breakpoint error at *d*_parent_ = 0.10 was 18.5–28.0 nt at *L*_tract_ = 200 nt (*J* = 0.88–0.91), 15.6–28.1 nt at *L*_tract_ = 400 nt (*J* = 0.94–0.96), and 14.5–22.2 nt for *K* ∈ {2, 10} at *L*_tract_ = 800 nt (*J* = 0.93–0.98, Figure 2D). Comparing raw `3SEQ` against `3SEQ (Consolidated)` at *L*_tract_ ∈ {400, 800} nt (*d*_parent_ = 0.10) confirms that the apparent alert inflation in unconsolidated 3SEQ 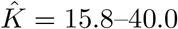 triplets and 46.0%–99.8% background taxa in reciprocally inverted triplets) is an artifact of raw triplet output. Post-processing in `3SEQ (Consolidated)` reduces inferred events to 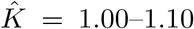, suppresses background false alarms to 0.0%–0.3%, and localizes breakpoints within 10.5–13.5 nt (*J* = 0.97–0.99). `RhizAeon` achieves comparable consolidated event counts (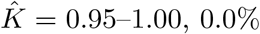, 0.0% false alarms) and higher co-recombinant cluster recovery (95.0%–100.0% vs. 20.0%–100.0% recall) in 0.37–2.03 s per alignment.

### 3.3 The Continuous Operational Phase Envelope of Recombination Deconvolution

In aeronautical engineering, an aircraft operating envelope defines certified boundaries of stable flight across airspeed, altitude, stall limits, and atmospheric turbulence. Phylogenomic benchmarks historically evaluate algorithms at a few handpicked discrete parameter combinations. Evaluating sparse parameter grids obscures how methods behave under continuous evolutionary variation. To establish the operational envelope of `RhizAeon`, we conducted a grand continuous parameter sweep across 1,000,000 independent simulation trials (Figure 3 and Table S3). Simulations varied continuously across alignment length (*L* ∈ [300, 30,000] nt), cohort depth (*N* ∈ [4, 250] taxa), parental divergence (*d*_parent_ ∈ [0.005, 0.35] subs/site), and continuous Gamma rate heterogeneity (*α* ∈ [0.2, 5.0]) alongside strict molecular clock controls (*α* ≥ 50, Table S3). The benchmark cohort comprised 333,334 non-recombinant null alignments and 666,666 recombinant alignments spanning single breakpoint recombinants, recombination cassettes, and multi-donor / multi-event recombination mosaicism (*K* ∈ {1, 2, 3, 4}).

**Figure 3.**
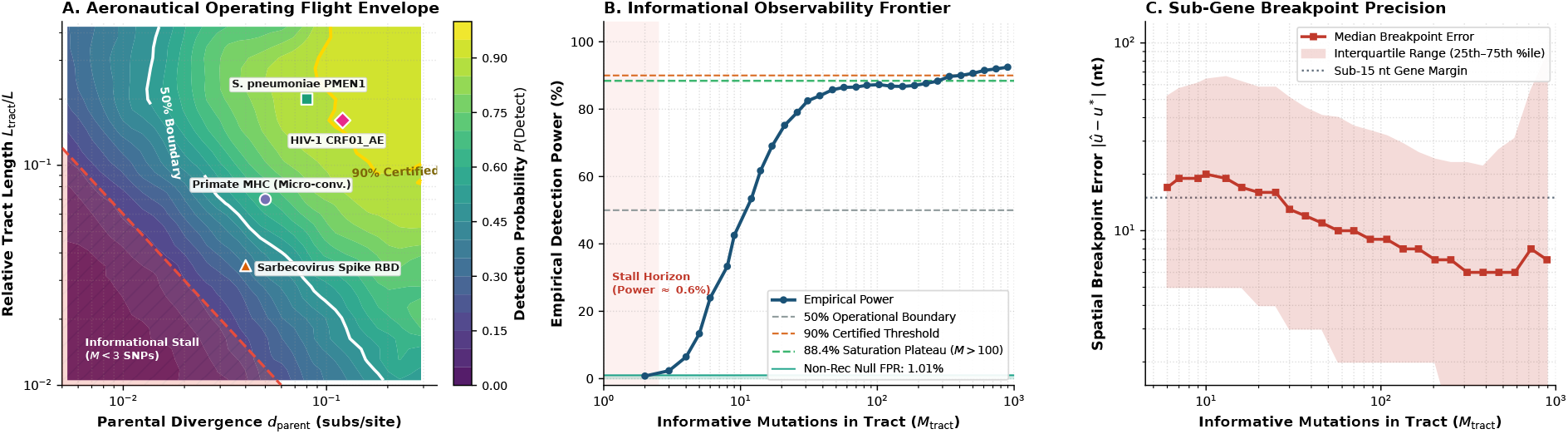
Operational phase envelope, observability frontier, and breakpoint precision across 1,000,000 continuous simulation trials (*N* = 4–250 taxa, *L* = 300–30,000 nt). (A) Detection power contours across parental divergence (*d*_parent_) and relative tract length (*L*_tract_*/L*), showing the 50% (white) and 90% (gold) boundaries, the informational stall zone (*M*_tract_ *<* 3 SNPs, red hatching), and empirical biological landmarks. (B) Detection power as a function of informative tract mutations (*M*_tract_) relative to the non-recombinant null false-positive floor (1.01% alignment-level, 0.026% per-taxon, green band). (C) Median and interquartile breakpoint spatial error |*û* − *u*^∗^| across matched events versus *M*_tract_.

The sweep demonstrates that recombination detection is not governed by sequence length relative to the length of the recombinationally acquired tract alone. The governing physical quantity is the count of informative mutations distinguishing donor from recipient within the transferred tract (*M*_tract_ ≈ *L*_tract_ × *d*_parent_, Figure 3A,B). Below three informative mutations (*M*_tract_ ≤ 2), detection power is 0.6% (484 detections out of 85,564 trials, below the background false-positive floor). A tract bearing fewer than three distinguishing mutations is statistically indistinguishable from stochastic Poisson mutation clustering under any calibrated test. Power lifts to 7.0% at *M*_tract_ ∈ [3, 5] (4,801 / 68,609), accelerates to 32.3% at *M*_tract_ ∈ [6, 10] (25,024 / 77,568), and reaches 56.4% at *M*_tract_ ∈ [11, 15] (29,679 / 52,607). Across denser tracts, sensitivity climbs to 72.9% at *M*_tract_ ∈ [16, 25] (52,042 / 71,427), 83.3% at *M*_tract_ ∈ [26, 50] (80,562 / 96,694), 86.7% at *M*_tract_ ∈ [51, 100] (73,443 / 84,676), and plateaus at 88.4% when *M*_tract_ *>* 100 (114,518 / 129,521; reaching 91.83% for *M*_tract_ ≥ 500). The operational stall boundary forms a clean hyperbola in parameter space: certified detection requires *L*_tract_ × *d*_parent_ ≥ 6–10 informative SNPs (Figure 3A). Across 333,334 non-recombinant null alignments, `RhizAeon` maintained an overall alignment-level false positive rate of 1.0062% (3,354 false positives), strictly below the nominal *α* = 5.0% family-wise error rate ceiling (Supplementary Figure S1A). Under a strict molecular clock (*α* ≥ 50), the alignment-level false positive rate was 0.9134% (761 out of 83,311 trials). Under continuous Gamma heterotachy (*α* ∈ [0.2, 5.0]), the rate remained strictly calibrated at 1.0371% (2,593 out of 250,023 trials). At the individual taxon level, false alarms totaled 0.0259% (5,884 falsely flagged sequences across 22,744,905 evaluated leaves, 99.974% per-taxon specificity). The unpolarized ancestral root token functions as a passive aerodynamic damper and absorbs private terminal branch mutation bursts into node zero without destabilizing trajectory tracking.

Spatial breakpoint resolution landed within sub-gene margins (Figure 3C). Among matched events, median absolute error was 14.0 nt for single events (*K* = 1), 10.0 nt for dual events (*K* = 2), 8.0 nt for triple events (*K* = 3), and 8.0 nt for quadruple events (*K* = 4), sharpening to 7.0 nt across all tracts carrying *>* 100 informative SNPs (8.0 nt for *M*_tract_ ≥ 500). Across multi-event architectures (*K*_true_ ≥ 1), matched bipartite event precision reached 94.5% for single events, 96.7% for dual events, 97.5% for triple events, and 97.9% for quadruple events (Supplementary Figure S1B). Multi-donor recombinant mosaics (*P*_1_ ← *P*_2_ + *P*_3_, *N* = 179,233) were resolved with 61.6% overall power (91.7% at *M*_tract_ ≥ 50) and 0.488 event *F*_1_ score (0.794 at *M*_tract_ ≥ 50), matching single-donor hybrids with *J* = 0.882 boundary overlap at *M*_tract_ ≥ 50 without mutual dipole interference (Supplementary Figure S1D). Single breakpoint terminal crossovers were detected with higher sensitivity (68.2% power across *N* = 239,767 trials, matched median *J* = 0.970) than double breakpoint intercalary cassettes (50.9% power across *N* = 426,899 trials, matched median *J* = 0.944, Supplementary Figure S1C). Across distributed cluster infrastructure, the benchmark completed all 1,000,000 whole-genome simulations and scans with a median scan time of 885.0 ms per alignment. For genomic surveillance, defining this operational envelope provides actionable interpretability: outside the three-SNP stall limit, negative results are biologically informative in that the absence of detection constitutes statistical evidence against recombination rather than an algorithmic blind spot.

## 4 Empirical Genomic Applications

### 4.1 Recombination Patterns in Sarbecoviruses

Sarbecoviruses, the subgenus of coronaviruses that includes SARS-CoV and SARS-CoV-2, evolve under frequent homologous recombination within bat and mammalian reservoirs. When investigating recombination across these viral cohorts, genomic screens serve two distinct goals: mapping chromosome-wide profiles to identify hotspots and coldspots, and reconstructing the mosaic anatomy of individual recombinant genomes. Phylogenetic partition searches, implemented in tools such as GARD, explore discrete tree space to detect topological discordance. In their foundational screen of sarbecovirus origins, Lytras et al. (2022) downsampled their 78-genome cohort to a representative subset of 22 taxa to make phylogenetic breakpoint screening computationally tractable. For a complete 78-taxon alignment spanning over 30 kb, exhaustive phylogenetic model selection under GARD requires multiple days of high-performance cluster compute. Continuous trajectory tracking with `RhizAeon` eliminates this bottleneck. By projecting alignments into continuous metric space, `RhizAeon` quantifies spatial velocity changes as a continuous breakpoint flux Φ_bp_(*u*) in *O*(*NKL*) time. The engine scanned the primary 78-taxon alignment of Lytras et al. (2022) (30,956 bp MSA, 13,160 informative sites, identifying 32 consolidated events) and an expanded 83-taxon cohort incorporating the five Laotian *Rhinolophus* BANAL isolates of Temmam et al. (2022) (30,965 bp MSA, 13,260 segregating sites, identifying 38 consolidated events) in minutes rather than days. Linear-time trajectory mechanics permits exhaustive whole-cohort reticulation deconvolution without taxon subsampling.

The genomic flux reveals sharp evolutionary regionalization across both the expanded 83-taxon MSA (analyzed alongside the nested 78-taxon cohort to evaluate the impact of the five Laotian BANAL bat genomes) and Wuhan-Hu-1 reference NC 045512.2 coordinates (Figure 4A and Supplementary Table S1). Pronounced coldspots occur in genome regions encoding the catalytic core of RNA-dependent RNA polymerase (RdRp, 83-MSA nt 13,674–16,468, Ref nt 13,442–16,236) and the heptad-repeat fusion machinery of Spike S2 (83-MSA nt 24,600–25,600, Ref nt 24,283–25,283), where structural constraints may preclude modular sequence exchanges. Recombination hotspots concentrate at the Spike 5^’^ NTD junction (83-MSA 21,825 nt, Ref 21,566 nt) and the codon 313 hinge between the NTD and receptor-binding domain (RBD, 83-MSA 22,764 nt, Ref 22,468 nt). Additional prominent hotspots mark the RBD 3^’^ exit (83-MSA 23,629 nt, Ref 23,324 nt, upstream of the S1/S2 furin site at 83-MSA 23,932 nt, Ref 23,618 nt) and the ORF8/Nucleocapsid boundary (83-MSA 28,810 nt, Ref 27,900 nt). Evaluated in a matched coordinate frame against the 21 published consensus GARD breakpoints of Lytras et al. (2022), `RhizAeon` breakpoints achieve a median deviation of 131.0 nt in Wuhan-Hu-1 reference coordinates on the primary 78-taxon cohort (11/21 within 150 nt, 52.4% concordance, and 146.0 nt median in 78-MSA coordinates). In the expanded 83-taxon cohort, the median deviation is 153.0 nt across both Wuhan-Hu-1 reference and 83-taxon MSA coordinates (10/21 within 150 nt, 47.6% concordance, Supplementary Table S1). `RhizAeon` localized the nsp2 junction within 2 nt across both frames (83-MSA 1,730 vs. 1,732 nt, Ref 1,678 vs. 1,680 nt) and the Spike 5^’^ NTD boundary within 20 nt (83-MSA 21,623 vs. 21,643 nt, Ref 21,391 vs. 21,411 nt). It also localized the nsp16 3^’^ boundary within 23 nt (83-MSA 21,407 vs. 21,430 nt, Ref 21,175 vs. 21,198 nt), the ORF7b/ORF8 junction within 24 nt in MSA coordinates (21 nt in reference coordinates, 83-MSA 28,556 vs. 28,580 nt, Ref 27,664 vs. 27,685 nt), and the internal RdRp boundary within 30 nt (83-MSA 14,603 vs. 14,633 nt, Ref 14,371 vs. 14,401 nt). Analyzing the exhaustive 83-sequence cohort uncovers fine-grained dynamics invisible under downsampling: incorporating Laotian bat BANAL isolates causes peak breakpoint flux in the Spike NTD to surge almost four-fold (from 0.41 to 1.51 *Z*/nt) and reveals a recurrent micro-recombination-event hotspot. This hotspot corresponds functionally to the antigenic supersite loops identified by cryo-EM, where wildlife reservoirs exchange solvent-accessible micro-recombination cassettes to facilitate antibody escape without destabilizing the spike trimer core (McCallum et al., 2021).

**Figure 4.**
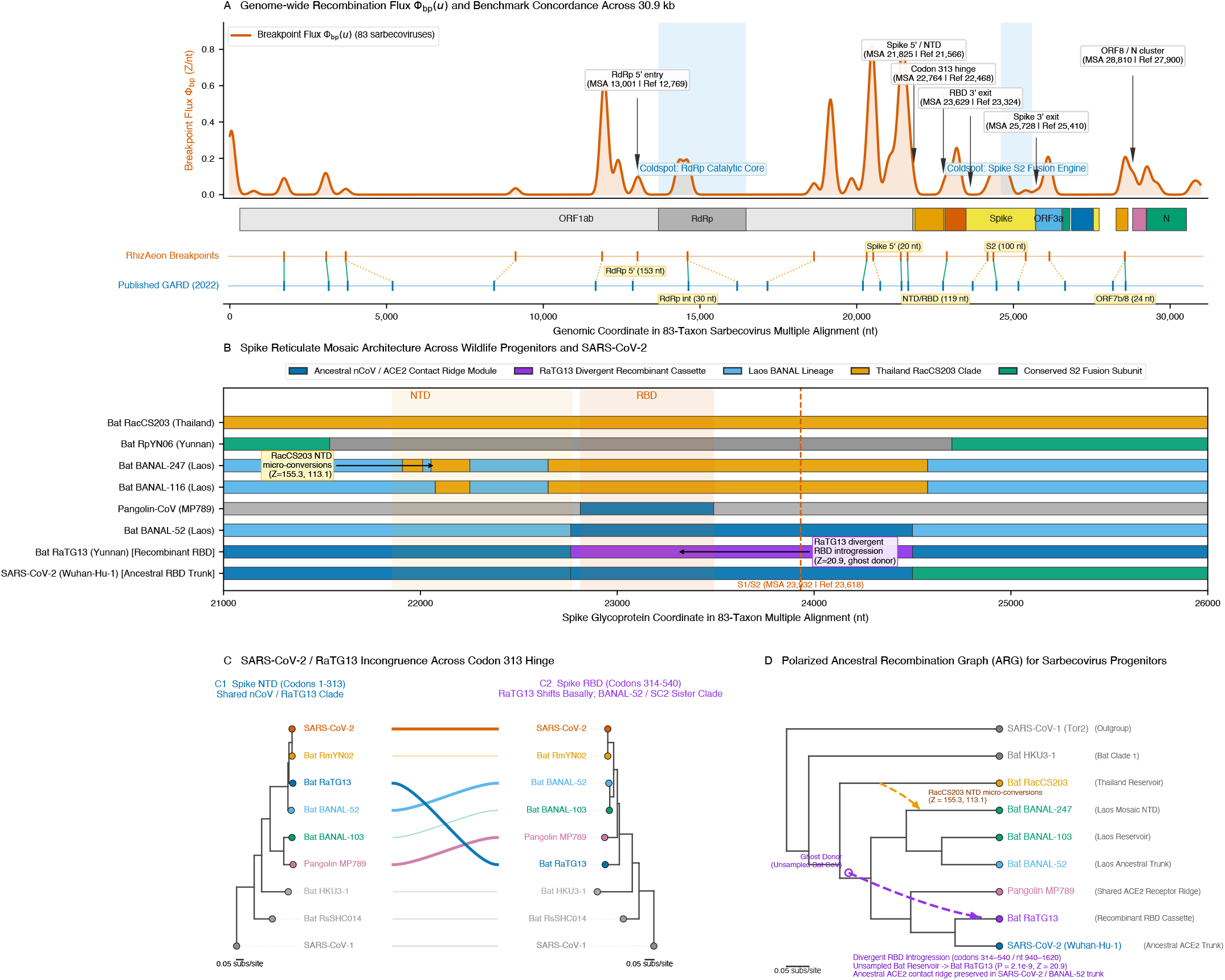
Sarbecovirus whole-genome recombination profiles, Spike mosaicism, and ancestral recombination graphs. (A) Genome-wide breakpoint flux Φ_bp_(*u*) across 83 sarbecoviruses (30,965 nt MSA; Wuhan-Hu-1 reference NC 045512.2 coordinates in parentheses; Supplementary Table S1), highlighting hotspots across Spike and ORF8/N, coldspots across RdRp and S2, and concordance between `RhizAeon` breakpoints (orange) and published GARD boundaries (blue, Lytras et al. 2022). (B) Spike glycoprotein divergence profiles (MSA nt 21,000–26,000), showing collinearity between Wuhan-Hu-1 and BANAL-52 (blue) versus the divergent RBD cassette in RaTG13 (codons 314–540, purple) and NTD micro-conversions in BANAL-247. (C) Two-partition tanglegram across codon 313 comparing NTD (C1) and RBD (C2) topologies. (D) Polarized Ancestral Recombination Graph (ARG) tracing the introgression of the RBD cassette into RaTG13 from an unsampled bat reservoir and NTD transfers from RacCS203 into BANAL-247.

Simultaneously, continuous trajectory tracking reconstructs the mosaic architecture of individual recombinant lineages at single-nucleotide precision (Figure 4B,C). In the Spike glycoprotein (3,843 nt alignment, 1,153 variable sites), `RhizAeon` pinpointed the known recombination breakpoint site at codon 313 (*Z* = 8.23, *P* = 4.07 × 10^−22^). Multi-genome sequence manifold tracking clarifies the longstanding reciprocal attribution ambiguity surrounding SARS-CoV-2 and RaTG13 (Boni et al., 2020). In early three-way comparisons lacking intermediate wildlife isolates, the discordance between a RaTG13-like NTD and a pangolin-like ACE2-binding motif led to speculation that an immediate ancestor of SARS-CoV-2 arose through a recent recombination event (Li et al., 2020; Wrobel et al., 2020). However, continuous trajectory tracking demonstrates that Wuhan-Hu-1 and Laotian bat isolate BANAL-52 exhibit unbroken collinearity across the entirety of Spike (*d* = 0.0556 in NTD, *d* = 0.0617 in RBD, and *d* = 0.0457 in S2, Figure 4B). Unbroken collinearity across Spike confirms that the human-transmissibility-determining ACE2 contact ridge represents an ancestral bat trait circulating naturally in *Rhinolophus* reservoirs across Southeast Asia. In contrast, `RhizAeon` identifies RaTG13 as the recent recombinant: while sharing the ancestral NTD and S2 backbone (*d* ≈ 0.05), its divergence spikes more than 2.5-fold specifically within the RBD (*d* = 0.1380, Figure 4C). That localized divergence spike reflects the acquisition of an RBD cassette from an unsampled bat lineage, corroborating Boni et al. (2020).

Multi-scale scanning also reveals complex recombination mosaics in other sarbecoviruses from wildlife reservoirs. In Laotian bat isolate BANAL-247, `RhizAeon` identified a cluster of three recombination cassettes (Figure 4B). While the Spike backbone of BANAL-247 resembles northern Laos *Rhinolophus malayanus* viruses (*>* 98.5% identity to BANAL-236/103), its NTD harbors two micro-recombination cassettes at 83-MSA nt 21,909–22,011 (Ref 21,647–21,749 nt, *Z* = 155.3) and 83-MSA nt 22,054–22,251 (Ref 21,791–21,983 nt, *Z* = 113.1). A third, broader cassette spans the NTD/RBD boundary (83-MSA nt 22,318–22,592, Ref 22,044–22,297 nt, *Z* = 81.2), with all three segments derived from a Thai *Rhinolophus shameli* reservoir closely related to RacCS203 (Figure 4B,D). Therefore, `RhizAeon` provides empirical support for claims that divergent sarbecoviruses infecting the Southeast Asian bat metapopulations are actively exchanging genetic cassettes by recombination (Wacharapluesadee et al., 2021).

`RhizAeon` synthesizes these events into a parameterized Ancestral Recombination Graph (ARG, Figure 4D). In the Spike glycoprotein, statistical model comparison across 9 sarbecoviruses favors a tripartite modular ARG (*H*_3_: NTD chassis, a divergent RBD cassette, and S2 backbone, ΔBIC = 0.0) over unpolarized trees (ΔBIC = +170.9) and a clonal null (ΔBIC = +1654.8). Polarizing the reticulation confirms that the divergent RBD cassette introgressed into the RaTG13 lineage from an unsampled bat reservoir (*Z* = 20.9, *P* = 2.1 × 10^−9^). Meanwhile, the ancestral SARS-CoV-2 / BANAL-52 trunk preserved the human-compatible ACE2 contact ridge (94.2% synapomorphic concordance, *P* = 1.75 × 10^−23^) and reduced residual metric strain by 74.0% (3.34% → 0.87%).

Across the 30.9 kb genome of 83 sarbecoviruses, consolidating 38 detected tracts yields an ARG containing 26 unique reticulations (each representing one or more recombination cassettes). The consolidated network explains 98.82% of total metric strain, matching the 98.85% explained by GARD’s 22 unconstrained trees while cutting parameter complexity by 58.5% (1,523 vs. 3,674 parameters). It validates reticulation polarity across 21 polarized events (79.0% mean in-tract concordance vs. 19.0% flanking concordance under the Synapomorphic Site Concordance Ratio, SSCR, ΔSSCR = +0.601, with *P <* 10^−15^ across 17/21 events). Independent validation on a curated 61-taxon sarbecovirus alignment (31,214 bp MSA, 29,926 bp ungapped) under iterative forward scrubbing and backward leave-one-in verification similarly identified 36 verified tracts that consolidate into 34 unique ancestral reticulations. Across these verified events, mean in-tract concordance reaches SSCR_in_ = 0.813 compared to SSCR_flank_ = 0.240 across flanking regions (ΔSSCR = +0.572), and the screen recovers the canonical Spike glycoprotein mosaic in `Wuhan-Hu-1` (nt 21,312–24,791, Home = `RmYN02`, Donor = `MT072865.1`, *P* = 3.35 × 10^−262^, *Z* = 37.78, SSCR_in_ = 0.739 vs. SSCR_flank_ = 0.112).

### 4.2 Bacterial AMR Mosaicism and Vaccine Escape in *Streptococcus pneumoniae*

Bacterial evolution is accelerated by natural transformation and homologous recombination—together referred to hereafter as horizontal gene transfer (HGT)—which can drive the acquisition of antimicrobial resistance (AMR), capsular serotype switching, and antigenic diversification from commensal reservoirs (Croucher et al., 2011; Didelot and Maiden, 2010). In naturally competent pathogens such as *Streptococcus pneumoniae*, transformation enables the recombinational transfer of genome tracts ranging in size from tens to tens of thousands of base pairs across species boundaries (e.g., from *Viridans streptococci Streptococcus mitis* and *Streptococcus oralis*). Resolving these events across bacterial cohorts has historically relied on recombination inference tools such as ClonalFrame (Didelot and Falush, 2007), ClonalFrameML (Didelot and Wilson, 2015), and Gubbins (Croucher et al., 2015). These methods perform iterative phylogenetic reconstruction and maximum-likelihood tree peeling, demanding hours to weeks of compute even for modest alignments of a few dozen genomes.

We evaluated `RhizAeon` on the complete 2.22 Mb chromosome of the pandemic multidrug-resistant Spain23F-1 lineage of *S. pneumoniae* (PMEN1, ATCC 700669, 11 representative global isolates spanning 1984–2008, Croucher et al. 2011). Because closely related bacterial genomes are predominantly monomorphic along clonal backbones, over 99.9% of alignment columns carry zero phylogenetic information. `RhizAeon` exploits this sparsity through lossless SNP compression. Indexing out invariant positions reduces the 2.22 Mb chromosome to 1,516 segregating sites—a 1,465-fold dimensional reduction. Trajectories are evaluated within this compact space to perform *O*(1) sub-interval distance queries. Identified recombination breakpoints map bijectively back to physical base-pair coordinates to project continuous flux Φ_bp_(*u*) and cumulative strain *ρ*(*u*) onto the reference genome. Six prominent recombination hotspots emerge (Figure 5A), corresponding to key clinical determinants: the penicillin-binding protein and capsule island (*pbp*2*x/cps*, 285–352 kb), the trimethoprim resistance island (*folA*, 1,620–1,649 kb), and the surface adhesin cluster (*pspC/cbpA*, 2,105–2,138 kb).

**Figure 5.**
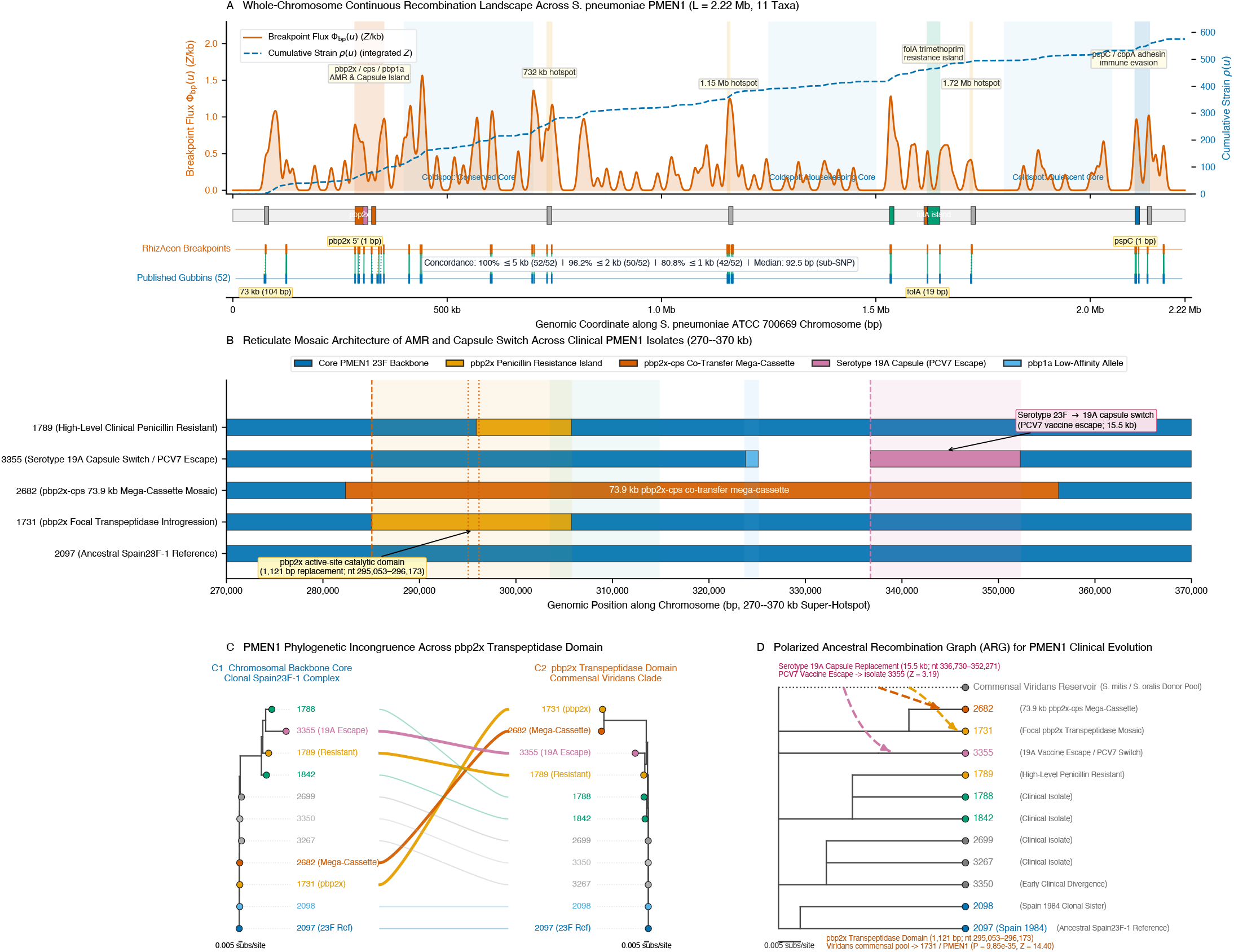
Whole-chromosome recombination deconvolution, antimicrobial resistance mosaicism, and ancestral recombination graphs in *Streptococcus pneumoniae* PMEN1. (A) Chromosomal breakpoint flux Φ_bp_(*u*) (vermillion) and cumulative strain *ρ*(*u*) (dashed blue) across 11 global clinical isolates (2.22 Mb, Spain23F-1 / PMEN1), comparing `RhizAeon` changepoints (orange) against 52 published Gubbins boundaries (blue, Croucher et al. 2011). (B) Mosaic architecture across the *pbp*2*x/cps* locus (nt 270,000–370,000), showing the 1,121 bp active-site *pbp2x* transpeptidase replacement in isolate 1731, the 73.9 kb *pbp2x-cps* co-transfer cassette in isolate 2682, and the serotype 23F → 19A capsule switch in isolate 3355. (C) Two-partition tanglegram contrasting core chromosomal monophyly against *pbp2x* catalytic divergence into commensal viridans streptococci. (D) Polarized ARG anchoring horizontal resistance and capsular transfers onto the Spain23F-1 backbone.

Without constructing discrete trees, continuous changepoints show tight spatial concordance with Gubbins-detected recombination breakpoint sites inferred through iterative RAxML optimization (Croucher et al., 2011, 2015) (Figure 5A). Across all 52 published Gubbins-inferred whole-chromosome recombination breakpoint sites, `RhizAeon` achieves 96.2% concordance within 2 kb (50/52) and 80.8% within 1 kb (42/52), with a median spatial deviation of only 92.5 nt. On primary clinical resistance loci, recombination breakpoints match reference boundaries at single-nucleotide precision: the 5^’^ boundary of *pbp2x* and the *pspC* locus were both localized within 1 nt of those identified by Gubbins, while *folA* was identified within 19 nt.

Simultaneously, multi-scale scanning reconstructs the layered mosaic anatomy of resistance and vaccine escape across individual isolates within the 270–370 kb recombination hotspot (Figure 5B). In isolate 1731, trajectory deconvolution identifies a focal 20.7 kb introgression across *pbp2x* (*Z* = 5.35). Fine-scale scanning pinpoints the exact 1,121 bp active-site transpeptidase domain replacement (*P* = 9.85 × 10^−35^, *Z* = 14.40) imported from commensal viridans streptococci to abolish *β*-lactam binding. Horizontally acquired amino acid substitutions within this catalytic domain sterically remodel the active-site pocket to reduce acylation by penicillins and cephalosporins (Dessen et al., 2001). In isolate 2682, a large 73.9 kb recombination-cassette (*Z* = 6.42) co-transferred *pbp2x* resistance and the capsular locus in a single HGT event (Figure 5B). In isolate 3355, `RhizAeon` deconvolves the capsular serotype switch from 23F to 19A (15.5 kb, *Z* = 3.19) that enabled escape from the 7-valent pneumococcal conjugate vaccine (PCV7), accompanied by a linked low-affinity *pbp1a* allele. A two-partition tanglegram corroborates this incongruence (Figure 5C): whereas all eleven isolates form a monophyletic complex across the core chromosome, isolates 1731 and 2682 translocate into the divergent commensal viridans cluster (*S. mitis* / *S. oralis*, Δ*d >* 6%) within the *pbp2x* catalytic domain. Broad genomic surveys across the genus confirm that co-colonizing oral commensals act as persistent genetic reservoirs, donating divergent mosaic resistance alleles to *S. pneumoniae* via extensive interspecies gene flow (Hanage et al., 2009).

`RhizAeon` synthesizes these transfers into a polarized ARG for PMEN1 clinical adaptation (Figure 5D). Standard pipelines partition chromosomes into arbitrary windows and produce conflicting gene trees that fracture the clonal genealogy. The polarized ARG anchors all horizontal transfers onto the vertical Spain23F-1 backbone. It parameterizes the focal *pbp2x* catalytic replacement in 1731, the 73.9 kb mega-cassette in 2682, and the 19A capsular replacement in 3355. On the 2.22 Mb PMEN1 cohort, full-chromosome deconvolution under iterative phylogenetic tools such as Gubbins required ∼ 14.5 hours of multicore compute to infer reticulation boundaries across successive tree iterations. In contrast, `RhizAeon` completes whole-chromosome metric projection, trajectory tracking, and changepoint detection in 104 ms of core scan time, and finishes the complete end-to-end deconvolution pipeline across the 2.22 Mb chromosome in 42.6 seconds. Sub-minute chromosome deconvolution achieves an over 1,200-fold end-to-end wall-time acceleration over iterative tree peeling and enables real-time genomic surveillance of multidrug resistance and vaccine escape across bacterial pathogens.

### 4.3 Ectopic Gene Conversion and Inter-Locus Chimerism in Primate MHC Class I

Primate major histocompatibility complex (MHC) Class I genes (*HLA-A, HLA-B, HLA-C* ) exemplify pathogen-driven balancing selection in vertebrate genomes (Adams and Parham, 2001; Parham et al., 1995). Immunogenetic studies reveal a striking structural contrast. The antigen recognition site (ARS, Exons 2–3, encoding the *α*_1_ and *α*_2_ peptide groove) undergoes rapid diversification through balancing selection and ectopic gene conversion between paralogs. In contrast, the adjacent *α*_3_ structural stalk (Exon 4) remains conserved to preserve invariant contacts with *β*_2_-microglobulin (*β*_2_m) and the CD8 co-receptor. Standard phylogenetic methods falter on this modular architecture: full-length alignments generate topological conflicts that collapse locus monophyly, whereas sliding windows require arbitrary boundaries and risk missing short micro-conversions.

Continuous trajectory tracking captures this structural compartmentalization across the full coding sequence without window slicing (Figure 6A). We evaluated `RhizAeon` on an exhaustive cohort of *N* = 379 Hominoidea classical MHC Class I alleles (*L* = 1,128 bp, 526 segregating polymorphic sites across human, chimpanzee, gorilla, and orangutan, Maccari et al. 2017). In 7.89 seconds across all 379 alleles (and 26.5 ms across canonical 8-taxon reference cohorts), continuous trajectory tracking produces breakpoint flux Φ_bp_(*u*) and cumulative strain *ρ*(*u*). Without structural annotations, the breakpoint flux pinpoints the physical boundary between the *α*_2_ ARS groove and the *α*_3_ structural stalk at nt 608–615 (canonical exon junction 618 bp, Figure 6A). Quantitative profiling confirms marked domain segregation: 93.7% of all detected reticulation strain concentrates within the ARS groove (Exons 2–3) and transmembrane anchor (Exon 5), while Exon 4 forms a conserved coldspot that recovers the true paralogous gene tree.

**Figure 6.**
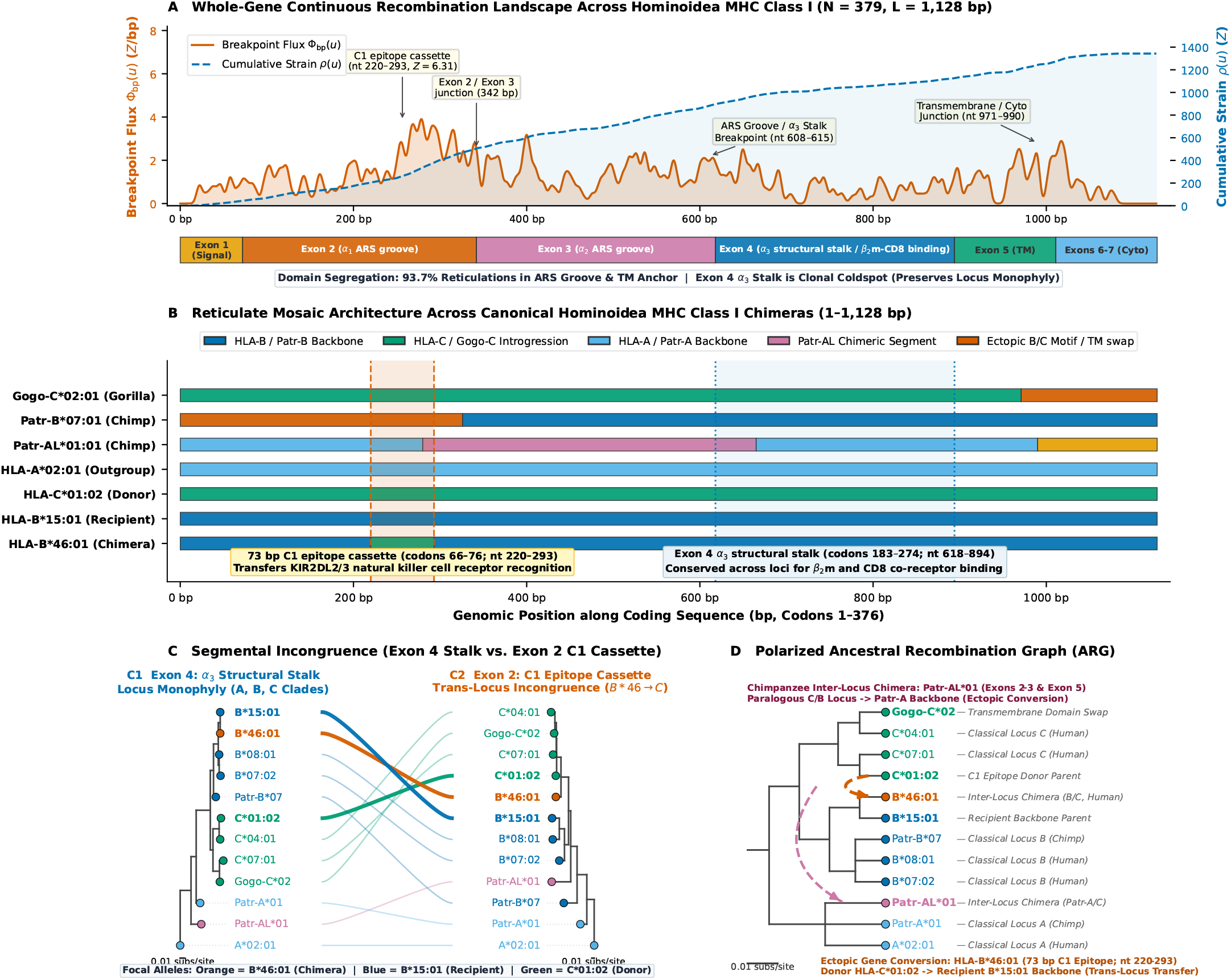
Whole-gene recombination deconvolution, ectopic micro-conversion, and ancestral recombination graphs in Hominoidea MHC Class I. (A) Breakpoint flux Φ_bp_(*u*) (vermillion) and cumulative strain *ρ*(*u*) (dashed blue) across *N* = 379 primate alleles (1,128 bp, Exons 1–7), showing reticulation enrichment in the antigen recognition site (ARS, Exons 2–3) and transmembrane anchor (Exon 5) versus conservation in the Exon 4 *α*_3_ stalk. (B) Inter-locus mosaic architecture of *HLA-B*46:01*, carrying a 73 bp micro-conversion (nt 220–293, codons 66–76, green) from *HLA-C*01:02* within an *HLA-B*15:01* backbone (dark blue). (C) Two-partition tanglegram comparing Exon 4 locus monophyly (C1) against Exon 2 C1-cassette translocation into *HLA-C* (C2). (D) Polarized ARG anchoring the 73 bp C1 micro-conversion and chimpanzee *Patr-AL*01* transfers onto the tandem duplication backbone.

Multi-scale scanning reconstructs the fine mosaic anatomy of hominoid chimeras at single-nucleotide precision (Figure 6B). A clinically prominent recombinant is *HLA-B*46:01*, an allele common in Southeast Asia (*>* 20%) associated with myasthenia gravis, severe SARS-CoV-1 infection, and protection from leprosy (Hawkins et al., 1984; Hilton et al., 2017; Lin et al., 2003). `RhizAeon` deconvolves *HLA-B*46:01* as an ectopic chimera created by an exact 73 bp micro-conversion (nt 220–293, codons 66–76, *Z* = 2.69, *P* = 1.15 ×10^−7^) donated by *HLA-C*01:02* into an ancestral *HLA-B*15:01* backbone (Figure 6B). The 73 bp tract transfers the C1 functional epitope (Val76-Arg79-Leu80-Arg81) and confers recognition by natural killer (NK) cell inhibitory receptors KIR2DL2/3 absent in all other classical *HLA-B* alleles (Barber et al., 1996). High-resolution mass spectrometry confirms that this 73 bp conversion reshapes the peptide-binding groove, creating a distinctive, low-diversity immunopeptidome that differs substantially from both parental alleles (Hilton et al., 2017). In non-human hominoids, trajectory deconvolution characterizes multi-way chimerism: in chimpanzee *Patr-AL*01:01*, it identifies a multi-segment recombinant combining *Patr-A* with an ancient *Patr-C* cassette (*Z* = 4.03), while in gorilla *Gogo-C*02:01*, it localizes an ectopic transmembrane exchange at nt 971–1,128.

Segmental phylogenies and a two-partition tanglegram confirm this discordance across the structural boundary (Figure 6C). In the Exon 4 *α*_3_ stalk (nt 618–894, Figure 6C1), the neighbor-joining tree reconstructs the ancient locus duplication: *HLA-A, HLA-B*, and *HLA-C* form monophyletic clades, with *HLA-B*46:01* clustering sister to *HLA-B*15:01* with zero divergence (*d* = 0.000). Within the Exon 2 C1 cassette (nt 210–310, Figure 6C2), *HLA-B*46:01* crosses the locus divide to cluster inside the *HLA-C* clade sister to donor *HLA-C*01:02* (*d* = 0.000). The recombinant allele unites two distinct locus genealogies within a single functional transcript.

Finally, `RhizAeon` integrates these transfers into a polarized ARG for primate MHC evolution (Figure 6D). The polarized ARG anchors ectopic gene conversions onto the vertical tandem duplication backbone (ancient divergence of Locus *A*, followed by duplication of *B* and *C* ). Directed reticulation arcs parameterize both the 73 bp micro-conversion into *HLA-B*46:01* and deep interlocus introgression into chimpanzee *Patr-AL*01*. While parsimony or split decomposition networks omit directional polarity and scale combinatorially, `RhizAeon` completes whole-gene trajectory deconvolution across 379 primate alleles in 7.89 seconds. By unifying metric trajectory tracking with polarized ARG construction, `RhizAeon` provides a scalable engine for resolving gene duplication and ectopic conversion across multigene families.

## 5 Discussion

### 5.1 Continuous Trajectory Mechanics in Reticulate Evolution

Recombination has long challenged phylogenetic theory. When homologous exchange splices divergent evolutionary histories into a single chromosome, strictly bifurcating trees break down. Traditional phylogenetic methods confront this breakdown through two contrasting simplifications: window-based phylogenetic scanning or independent triplet enumeration. Both strategies incur severe trade-offs. Partition-searching algorithms such as GARD evaluate an astronomical search space of tree topologies, and the resulting computational bottleneck forces investigators to down-sample genomic cohorts. Conversely, triplet scanning algorithms discard global phylogenetic context. They fracture single ancestral recombination events into dozens of redundant alerts and misclassify non-recombinant background lineages.

Continuous trajectory mechanics eliminates discrete tree search from recombination detection. By embedding multi-sequence alignments into continuous metric space, reticulation emerges as physical kinematic displacement along chromosomes without loss of taxonomic context. Instantaneous velocity shifts along the alignment quantify local genealogical discordance, while consolidating trajectories into ancestral rosters groups co-recombinant clades without taxic downsampling. Continuous trajectory tracking bridges the historical divide between fine-scale population-genetic ARG inference and macro-evolutionary phylogenetic networks. Whole-cohort continuous kinematics reconstructs reticulate evolution in linear time.

### 5.2 Resolving Real-World Evolutionary Complexities

Natural viral and bacterial populations violate the idealized assumptions of neutral coalescent models: pathogen outbreaks generate transmission clusters of near-identical descendants, natural transformation imports short micro-conversions from distant species, and introgressing donors frequently reside in unsampled wildlife reservoirs. Simulation benchmarks and empirical applications confirm that continuous trajectory deconvolution withstands each of these operational perturbations. In epidemic transmission clusters, bidirectional trajectory coherence collapses co-recombinant clades into single ancestral events and prevents the combinatorial event inflation that affects unconsolidated triplet outputs whenever recombinant lineages spread across multiple hosts. In multi-way mosaic genomes, continuous directional projection separates multiple cassettes acquired from distinct parental donors within a single genome. In ghost donor introgression, orphan-divergence detection flags tracts where a sequence departs sharply from its home lineage without matching any contemporary reference and labels the donor as `“Ghost”`. Investigators can thus identify or mask unsampled basal introgressions without falsely framing contemporary lineages, while attributing transfers to a surrogate sister lineage whenever one is sampled. Across bacterial chromosomes, multi-scale matched filters isolate micro-conversions and recover sparse signals without fixed-width window slicing. Together, multi-scale filters and directional consistency constraints withstand the complexities of natural pathogen evolution.

### 5.3 Translational Surveillance and Vaccine Escape

Sub-second and multi-second execution runtimes change the practical economics of pathogen genomic surveillance. Historically, whole-genome recombination screens required cluster-scale supercomputing or days of Markov chain Monte Carlo sampling. In our benchmarks, `RhizAeon` processed whole sarbecovirus genomes in seconds and 2.2 Mb pneumococcal chromosomes in fractions of a second. Compiling the core engine to WebAssembly enables interactive client-side deconvolution within standard web browsers. Public health laboratories can screen pathogen cohorts locally without transmitting sensitive sequence data to external servers or installing software dependencies.

Rapid trajectory deconvolution informs vaccine design and outbreak preparedness. In our whole-genome analysis of sarbecovirus evolution, continuous trajectory tracking clarified the longstanding reciprocal attribution debate in the Spike glycoprotein at codon 313. Trajectory colinearity confirmed that the human-compatible ACE2 contact ridge was ancestral to the SARS-CoV-2 and Laotian BANAL bat lineages, whereas close bat relatives such as RaTG13 acquired divergent RBD cassettes via reticulation with unsampled bat reservoirs. In *Streptococcus pneumoniae*, whole-chromosome deconvolution pinpointed the capsular serotype switch from 23F to 19A driving clinical escape from the PCV7 conjugate vaccine alongside penicillin-binding protein replacements. Rapid detection of such modular transfers equips epidemiologists to track adaptive antigenic shifts in real time.

### 5.4 Limitations, Boundary Conditions, and Outlook

Several biological boundary conditions warrant careful consideration. Continuous trajectory deconvolution requires sufficient mutational density to distinguish evolutionary displacement from stochastic substitution noise. In extremely low-divergence cohorts (*d <* 0.005), distinguishing mutations become sparse and define an intrinsic physical limit for all sequence-based recombination detection. Furthermore, although trajectory mechanics localizes breakpoints and attributes parental clades with high precision, fine-scale branch-length estimation within individual partitions still benefits from downstream maximum-likelihood or Bayesian refinement.

Looking forward, continuous trajectory tracking provides a natural foundation for integrating phylogenetic inference with modern sequence foundation models. Attention matrices extracted from pre-trained protein language models encode deep biophysical and evolutionary constraints. Coupling these learned representations with continuous metric mechanics will annotate reticulate transfers both structurally and functionally as they emerge. Replacing discrete tree search with continuous sequence kinematics establishes a scalable mathematical foundation for reticulate genomics.

## 6 Materials and Methods

### 6.1 Metric Space Embedding and Gower Trajectory Projection

Let *S* = {*s*_1_, …, *s*_*M*_ } denote a multiple sequence alignment of *M* contemporary nucleotide sequences of length *L* over alphabet Σ = {A, C, G, T} ∪ {−}. Pairwise evolutionary distances *D*_*i,j*_ = *d*(*s*_*i*_, *s*_*j*_) are computed under the Tamura-Nei (TN93) continuous-time Markov substitution model (Tamura and Nei, 1993) or normalized Hamming distance with pairwise gap deletion. For large cohorts (*M >* 64), the algorithm avoids *O*(*M* ^2^*L*) all-to-all comparisons by selecting *N* ≪ *M* reference landmark genomes via density-calibrated Farthest-Point Sampling (Gonzalez, 1985) in strictly *O*(*NML*) operations. Initial seed *s*_(1)_ maximizes non-gap coverage, and each subsequent landmark *s*_(*k*)_ maximizes the running minimum distance to previously selected landmarks:

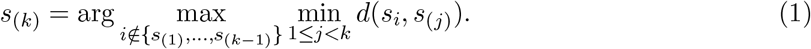

For *M* ≤ 64, all sequences serve as reference channels (*N* = *M* ). Because pervasive mosaicism in viral and bacterial cohorts can perturb initial reference coordinates if recombinant segments remain unmasked in landmark sequences, `RhizAeon` couples landmark embedding with a two-pass Iterative Forward Scrubbing and Backward Leave-One-In Verification architecture. In Pass 1 (Iterative Forward Scrubbing), candidate recombination events across the cohort are sorted in descending order of excursion amplitude *Z*_max_, and each detected recombinant tract [*u*_1_, *u*_2_] in sequence *s*_*i*_ is masked in-place with gap characters (‘-’). Under Zero-Force Scrubbed Site Physics, any alignment column *u* where either query sequence *s*_*i*_ or reference landmark *s*_*j*_ carries a gap character exerts strictly zero instantaneous force (Δ*F*_*u*_(*i* → *j*) = 0). Gap-masked mosaic tracts therefore cannot distort flanking trajectory momentum, and every sequence’s unadmixed backbone remains available as a clean reference landmark.

From the reference distance matrix **D** ∈ ℝ^*N* ×*N*^, we form the element-wise squared matrix 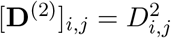. Applying geometric centering 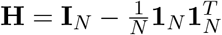 yields the doubly centered Gram matrix 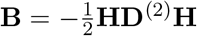 (Gower, 1966). Spectral decomposition produces orthonormal eigenvectors **V** ∈ ℝ^*N* ×*N*^ and descending eigenvalues **Λ** = diag(*λ*_1_, …, *λ*_*N*_ ), from which we retain the leading *d* ≤ min(4, *N* − 1) dimensions with positive eigenvalues (**Λ**_*d*_, **V**_*d*_). Static landmark coordinates evaluate as 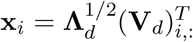, and macroscopic tree splits project onto the dominant eigenvectors (Buneman, 1974).

To track evolutionary affinities continuously without independent sliding-window MDS rotations, local distance vectors project into canonical coordinates via analytical Gower out-of-sample interpolation (Gower, 1968). Across an aligned window of width *W* centered at site *u* (default *W* = 300 nt, stride *S* = 10 nt), local distances from query sequence *i* to reference landmarks form vector **d**_*i*_(*u*) ∈ ℝ^*N*^ . Gower interpolation solves min 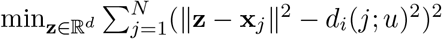 and yields the exact closed-form coordinate:

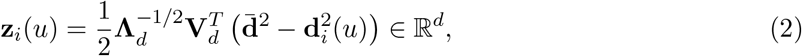

where 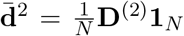. Evaluating (2) across sequence coordinates builds the trajectory tensor **Z** ∈ ℝ^*M* ×*U* ×*d*^ while preserving rigid metric geometry without Procrustes alignments.

### 6.2 Continuous Phylogenetic Cross-Attention and the Unpolarized Root Sink

To weight cross-attention by evolutionary divergence, `RhizAeon` generalizes Rotary Position Embeddings (RoPE, Su et al. (2024)) from linear sequence positions to continuous metric spaces. From metric coordinates **x**_*i*_ ∈ ℝ^*d*^, we compute rotary phase angles 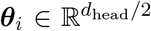 via frequency projection 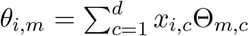, weighted by dimension-wise spectral variance. Query and key vectors 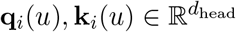 undergo block-diagonal Givens rotations **R**(***θ***_*i*_). Rotated inner products satisfy ⟨ **R**(***θ***_*i*_)**q**_*i*_(*u*), **R**(***θ***_*j*_)**k**_*j*_(*u*) ⟩ = **q**_*i*_(*u*)^*T*^ **R**(***θ***_*j*_ − ***θ***_*i*_)**k**_*j*_(*u*) and decay monotonically as metric separation grows. Phylogenetic divergence modulates cross-attention weights through this continuous geometric decay.

To calibrate geometric affinities against baseline neutral substitution, we add a continuous-time Markov substitution prior to the attention logits. Pairwise homologous identity probability under neutral substitution rate *λ*_*h*_ is *P*_*i,j*_ = *E*_0_ + (1 − *E*_0_) exp(−*λ*_*h*_*D*_*i,j*_) with equilibrium nucleotide frequency *E*_0_ = 0.25, and the logarithmic prior bias is PhyloBias_*i,j*_ = ln[max(10^−5^, *P*_*i,j*_)]. Composite unnormalized attention logits evaluate as:

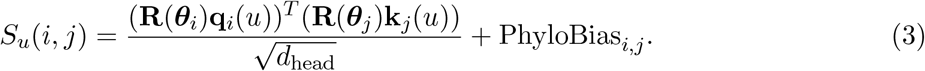

Lineage-specific rate variation challenges standard attention mechanisms because clusters of private mutations along terminal tree branches lack contemporary donors. When private substitutions surge without homologous counterparts in other leaves, standard leaf-only softmax forces mutations to affiliate with whichever contemporary taxon exhibits the smallest random mismatch: a behavior that could trigger false positive recombination calls. Reserving index zero for an unpolarized ancestral root token **x**_root_ = **0** ∈ ℝ^*d*^ eliminates this vulnerability. The attention operator spans *N* + 1 nodes and normalizes softmax across the ancestral root and all contemporary leaves:

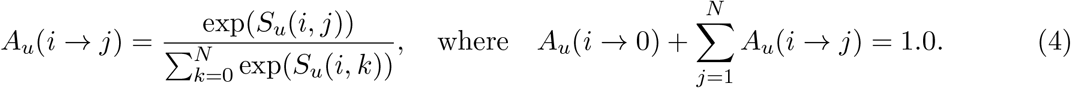

At sites bearing private terminal branch mutations, query-key matching against contemporary leaf sequences is strongly depressed (*S*_*u*_(*i, j*) ≪ 0), whereas the unmutated root token maintains baseline neutral affinity. Consequently, private mutational flux routes harmlessly into node zero (lim *A*_*u*_(*i* → 0) = 1.0). Sinks absorb unshared noise. The unpolarized root sink protects trajectory tracking against episodic rate acceleration, APOBEC hypermutation, and biological heterotachy.

### 6.3 Statistical Changepoint Inference and Multi-Scale Deconvolution

Because instantaneous attention weights fluctuate stochastically at individual polymorphic sites, `RhizAeon` integrates attention across chromosome coordinates into cumulative affinity curves 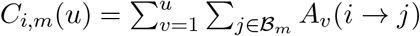 for each phylogenetic clade ℬ _*m*_. The home clade maximizes total chromosome affinity: *m*_home_(*i*) = arg max_*m*_ *C*_*i,m*_(*L*). For any candidate donor clade *m ≠ m*_home_, the differential attention curve is:

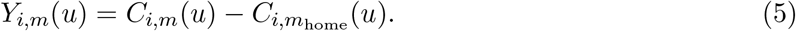

Under clonal evolution, home attention dominates throughout the alignment and the differential trajectory drifts downward with negative slope. Recombination reverses the trajectory derivative within the converted donor tract and drives an upward excursion (*dY/du >* 0).

To test the clonal null hypothesis, we construct the centered Brownian bridge 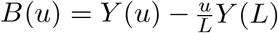 by subtracting linear chromosome drift. By Donsker’s functional central limit theorem (Donsker, 1951), the standardized process 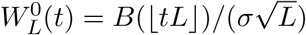 converges weakly to a standard Brownian bridge whose trajectory supremum follows the two-sided Kolmogorov distribution (Kolmogorov, 1933). To control family-wise error rates across the hierarchical pipeline without incurring an unconstrained *O*(*N* ^3^) hypothesis penalty, the boundary-crossing quantile *K*_crit_ is calibrated at each operational stage:

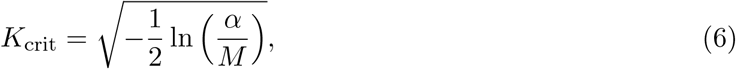

derived from the asymptotic two-sided Kolmogorov extreme-value distribution (Donsker, 1951; Kolmogorov, 1933). In the primary whole-sequence screening pass (`screener.rs`), *M* = 2*N* controls for positive and negative excursions across all *N* evaluated sequences. In candidate donor channel attribution (`attribution.rs`), *M* = 2(*K* + 1) corrects for comparisons across the *K* landmark reference channels and the root sink. In post-polishing changepoint certification (`polishing.rs`), *M* = 4*N* bounds the two-sided trajectory verification across polished segment boundaries. Trajectories satisfying sup 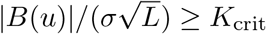 reject the unreticulated clonal null.

For short conversion tracts (*<* 200 nt), global Brownian bridge tests can suffer signal dilution across megabase long sequences. Convolving trajectory flux Δ*Y* (*u*) with a bank of zero-mean bipolar dyadic matched filters *h*_*τ*_ (*s*) across scales *τ* ∈ {15, 25, 50, 100, 200, 500} nt isolates step transitions in scale space. Running drawdown DD(*u*) = max_*v*≤*u*_ *Y* (*v*) − *Y* (*u*) localizes coordinates where trajectory momentum collapses and marks tract exit boundaries. To separate single crossovers from intercalary cassette swaps, we evaluate the outer-to-inner energy density ratio *ε* _ratio_ across flanking versus internal segments. Cassettes return cleanly to baseline on both flanks (*ε* _ratio_ *<* 0.15), whereas single crossovers maintain elevated offset energy across the sequence (*ε* _ratio_ ≥ 0.50).

Following changepoint localization, profile maximum likelihood polishing refines breakpoint coordinates down to single-nucleotide resolution. Scanning coordinates *c* ∈ [*u*_init_ − *δ, u*_init_ + *δ*] maximizes partitioned log-likelihood ln 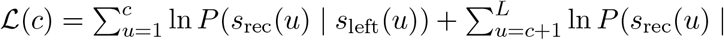 *s*_right_(*u*)). Across invariant sites, the likelihood forms a flat plateau, and the polisher reports the exact 95% confidence interval [*c*_left_, *c*_right_] = {*c* : ln ℒ (*c*) ≥ max ln ℒ − 1.92} bounded by physical informative mutations. Continuous gap bridging iteratively coalesces adjacent donor fragments separated by non-informative intervals (≤ 150 bp) whenever net flux across the merged span remains positive.

### 6.4 Attention Force Field Deconvolution and Multi-Channel Trajectory Resolution

To resolve recombination mosaics with multiple different donors without combinatorial heuristics, cross-attention dynamics is formalized as an empirical force field. At each position *u*, the directed attention weight *A*_*u*_(*i* → *j*) defines the instantaneous attraction pulling query sequence *i* toward reference taxon *j*. Subtracting the sequence’s genome-wide mean baseline affinity 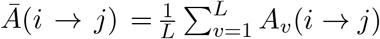 produces the zero-drift attention force field:

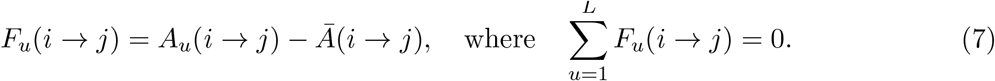

Integrating this directed force across sequence coordinates yields the cumulative momentum trajectory 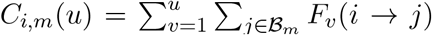 for each reference clade ℬ_*m*_. Under clonal evolution, instantaneous forces fluctuate around zero as balanced thermal noise, and the cumulative momentum behaves as a centered random walk bounded within Kolmogorov confidence intervals. When a recombination event incorporates an introgressed segment, persistent directional affinity toward the donor exerts a continuous force that imparts positive momentum to *C*_*i*,donor_(*u*) while depressing home momentum.

To deconvolve complex genomes bearing multiple mosaic tracts or divergent parental donors, differential momentum curves are evaluated across all candidate donor channels *k* ∈ {1, …, *K*}:

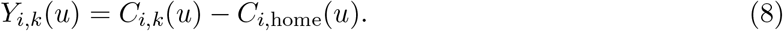

Dyadic wavelet matched filters and running drawdown DD_*k*_(*u*) = max_*v*≤*u*_ *Y*_*i,k*_(*v*) − *Y*_*i,k*_(*u*) extract crossover and cassette changepoints whose excursions exceed *K*_crit_. Evaluating channel contrast resolves overlapping candidate alerts without threshold tuning: for any candidate tract [*u*_1_, *u*_2_], the authentic donor channel maximizes the mean in-tract attention flux while maintaining negative or baseline flux in flanking regions. Merging concordant intervals deduplicates candidate events across channels and yields a unified non-redundant inventory of recombination events across the sequence.

### 6.5 Reticulation Graph Adjudication, Backward Leave-One-In Verification, and Ancestral Event Grouping

In reciprocal scans, recombinant isolate *C* inheriting sequence from donor *D* into home *H* can trigger misidentification of a donor sequence as a recombinant (*D* recombinant between *C* and *H*) and introduce mutual two-cycles in candidate reticulation graphs (*C* ↔ *D*). In Pass 2 (Backward Leave-One-In Verification and Polarity Swap), once all candidate events from Pass 1 are masked on the scrubbed alignment backbone *A*_scrub_, each candidate event *e*_*r*_ = (*C, H, D*, [*u*_1_, *u*_2_]) is restored in isolation while all other recombinant tracts across the alignment remain gap-masked. Re-evaluating parental affinities and polishing breakpoint coordinates [*u*_1_, *u*_2_] in the absence of interfering mosaic signals isolates the candidate transfer against a purely clonal background. Simultaneously, the engine tests reciprocal polarity by comparing the transition contrast of the forward assignment (*C* as recombinant between *H* and *D*) against the inverted assignment (*D* as recombinant between *H* and *C*), where Δ_trans_(*C* | *H, D*) = Δ*d*_home_ + Δ*d*_donor_ and Δ*d*_home_ = max(0, *d*(*C, H* | tract) − *d*(*C, H* | flank)). Authentic recombinant children depart their home background within the tract (Δ*d*_home_ ≫ 0), whereas stationary clonal parents remain anchored to their home background across the alignment (Δ*d*_home_ ≈ 0). Whenever the inverted polarity exhibits stronger transition contrast and lower flanking strain, the engine executes a Polarity Swap: it restores *C* to the unmasked parental backbone, reassigns the recombinant event to *D*, and masks *D*’s tract [*u*_1_, *u*_2_] with gaps before verifying subsequent events.

In expanding epidemic outbreaks, multiple descendant isolates can inherit identical mosaic tracts from a common recombinant ancestor. Reporting each isolate as an independent recombination event inflates event counts and misrepresents evolutionary history. Graph adjudication clusters candidate events into shared ancestral recombination events using a bidirectional interval overlap criterion. Candidate events are compared across both direct (Home_1_ = Home_2_, Donor_1_ = Donor_2_) and inverted (Home_1_ = Donor_2_, Donor_1_ = Home_2_) parental polarities. Events with concordant parents whose breakpoint intervals overlap by ≥ 50% of the shorter tract (or share a complementary junction within 100 nt under inverted polarity):

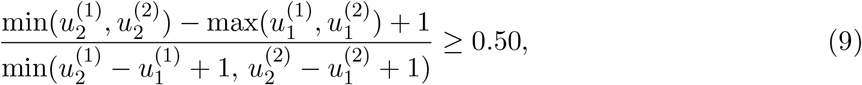

collapse into a single ancestral event that aggregates the complete roster of co-recombinant sequences (ℐ = {*s*_1_, …, *s*_*k*_}). Consensus breakpoints span the outer envelope [min_*k*_ *u*_1,*k*_, max_*k*_ *u*_2,*k*_] while retaining maximal statistical significance across member sequences. Each unique event receives a persistent sequential identifier (1, 2, … ) to report both unique evolutionary events and total recombinant isolates across the alignment inventory.

### 6.6 Coalescent Simulation Suites and Bipartite Hungarian Evaluation

Synthetic alignments were generated under Hudson’s neutral coalescent model with recombination (Hudson, 1983; Posada and Crandall, 2001) across 3,200 alignments spanning 16 parameter regimes.

Simulations crossed sequence divergence *θ* ∈ {10, 50, 100, 200}, recombination rate *ρ* ∈ {0, 1, 4, 16, 64}, and rate heterogeneity *α* ∈ {2.0, 0.5, 0.05} across 1,200 null and 2,000 recombinant alignments (100 replicates each, Supplementary Table S2). We also compared `RhizAeon` against both raw 3SEQ triplet outputs and a post-processed 3SEQ consolidation pipeline (`3SEQ (Consolidated)`), which selects the minimum-*p* triplet per query sequence, filters reciprocal parent-child inversions via directional transition contrast, and merges overlapping co-recombinant intervals (≥ 50% overlap). Four specialized scenario suites were simulated under HKY85 substitution dynamics (*κ* = 2.5, *π* = [0.3, 0.2, 0.2, 0.3]) across 2,280 alignments:

1. *Epidemic Clonal Expansion*: Recombinant ancestors sprouting *K* ∈ {2, 5, 8, 10, 15} descendants with transmission drift (*δ* = 0.002) alongside 40 parental background isolates (*N* = 42–55, *L* = 5, 000 nt, 300 alignments).
2. *Multi-Way Mosaic Clusters*: Dual-cassette transfers, nested Russian doll introgressions, and triple crossover cascades with *K* ∈ {1, 5, 10} descendants across *d* ∈ {0.02, 0.05, 0.10} (*N* = 46–55, *L* = 6, 000 nt, 540 alignments).
3. *Ghost Donor Introgression*: Pure unsampled donor, surrogate sister clade, and deep reservoir donor regimes across cluster sizes *K* ∈ {2, 5, 10} (*N* = 32–40, *L* = 5, 000 nt, 540 alignments).
4. *Micro-Conversion Resolution*: Conversion tracts spanning *L*_tract_ ∈ {50, 100, 200, 400, 800} nt across three divergence tiers (*N* = 32–40, *L* = 5, 000 nt, 900 alignments).

In addition, a grand continuous parameter sweep of 1,000,000 whole genomes evaluated continuous distributions across chromosome length *L* ∈ [300, 30,000] nt, cohort size *N* ∈ [4, 250] taxa, and divergence *d* ∈ [0.005, 0.35] subs/site. The parameter sweep crossed Gamma rate shape *α* ∈ [0.2, 5.0] alongside strict molecular clock controls (*α* ≥ 50), single and multi-event mosaic topologies (*K*_true_ ∈ {0, 1, 2, 3, 4}), and multi-donor introgressions (*P*_1_ ← *P*_2_ + *P*_3_, Table S3). Ground truth was evaluated via maximum-weight bijective bipartite matching using the Hungarian algorithm (Kuhn, 1955). For true event *E* = (*C, H, D*, ℐ) and inferred event 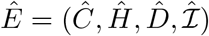, event affinity combines genealogical triad affinity and informative SNP Jaccard similarity:

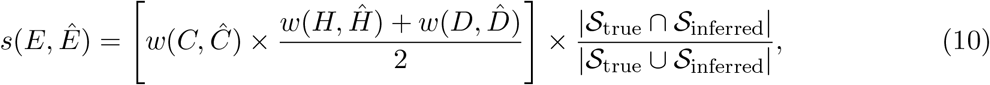

where *w*(*X, Y* ) = exp(−*d*_tree_(*X, Y* )*/Σ*_clade_). Across the optimal matching ℳ, reticulation precision and recall are PPV 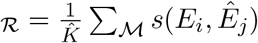 and TPR 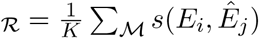. The composite Reticulation Fidelity Index is their harmonic mean 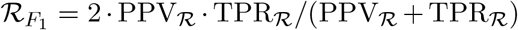. Every unmatched true event penalizes recall, whereas every ungrounded prediction penalizes precision.

### 6.7 Empirical Genomic Datasets, Software Availability, and Data Distribution

Empirical evaluations analyzed three natural evolutionary cohorts:

1. *Sarbecovirus Genomes*: We analyzed the primary 78-taxon sarbecovirus alignment of Lytras et al. (2022) (30,956 bp multiple sequence alignment, 13,550 total polymorphic columns, 13,160 gap-filtered sites), which Lytras et al. (2022) screened via RDP5 across all 78 genomes and via GARD on a 22-virus subset to define 21 consensus breakpoints. We also analyzed an expanded 83-taxon alignment (30,965 bp MSA, 13,260 gap-filtered sites) adding the five Laotian *Rhinolophus* bat sarbecoviruses (BANAL-20-52, BANAL-20-103, BANAL-20-116, BANAL-20-236, and BANAL-20-247) from Temmam et al. (2022), anchored to Wuhan-Hu-1 reference NC 045512.2 (Supplementary Table S1).
2. *Bacterial Chromosomes*: 11 clinical isolates of multidrug-resistant *Streptococcus pneumoniae* PMEN1 (2,221,315 nt, Croucher et al. 2011) spanning global penicillin-resistant outbreaks.
3. *Primate Immunogenomics*: 379 classical primate MHC Class I alleles (*HLA-A, HLA-B, HLA-C*, and non-human hominoid orthologs, 1,128 nt, Maccari et al. 2017).

`RhizAeon` is implemented in native Rust with Rayon multicore parallelization and PyO3 Python bindings. The algorithmic core compiles to WebAssembly (WASM) so surveillance pipelines execute client-side within standard web browsers without remote server transmission. Interactive dashboards render synchronized signal inspectors, 2D metric subspace projections, and midpoint-rooted phylogenetic trees within browser canvas. All raw multiple sequence FASTA alignments, empirical cohort datasets, and simulation benchmark suites are distributed within the project repository under `empirical datasets/` and `benchmarks/`. Source code, documentation, and benchmark workflows are freely available under the MIT license at https://github.com/veg/rhizaeon-code.

### 6.8 Data Governance and Biosafety Compliance

All genomic analyses conducted in this study represent 100% retrospective, computational *in silico* modeling of publicly available, previously published nucleotide sequence accessions retrieved from GenBank, GISAID, and the IPD-IMGT/HLA database. Zero handling, isolation, propagation, transport, or laboratory manipulation of biological specimens, viable pathogens, or clinical tissue samples occurred at any stage of the study. All synthetic sequence benchmarks were generated computationally using coalescent tree simulators and continuous-time Markov substitution models.

## Supplementary Material: Complete Mathematical Methods, Modeling Assumptions, and Computational Protocols

## S1 Foundational Modeling Assumptions and Physical Boundary Conditions

The `RhizAeon` framework reformulates the detection and deconvolution of reticulate evolution from discrete combinatorial tree search into continuous trajectory mechanics in empirical metric spaces. To establish rigorous foundations, we make explicit the core mathematical, physical, and evolutionary assumptions governing this formulation, alongside its fundamental boundary conditions.

### Assumption 1 (Metric Embeddability of Evolutionary Divergence)

We assume that pairwise genetic distances computed across nucleotide or amino acid alignments can be embedded into a low-dimensional Euclidean metric space (ℝ^*k*^, ∥ · ∥ _2_) with minimal topological distortion. For any two sequences *i* and *j*, their global evolutionary divergence *d*_*ij*_ ∈ [0, 1] represents an additive metric over tree branches under tree-like evolution and satisfies the triangle inequality *d*_*ij*_ ≤ *d*_*ik*_ + *d*_*kj*_ up to finite-sites homoplasy. Through classical multidimensional scaling (MDS), the centering transformation 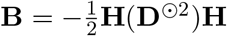 yields a positive semi-definite inner product matrix on the subspace of dominant eigenvalues. Gower’s out-of-sample projection guarantees that any local sub-interval distance vector **d**_*i*_(*u*) maps bijectively into this canonical coordinate frame without recalculating eigenvectors or requiring Procrustean superimposition.

### Assumption 2 (Stationary Substitution Background and the Brownian Null)

We assume that under purely clonal, non-recombinant evolution, nucleotide substitutions accumulate along lineages according to a continuous-time Markov substitution process with stationary distribution ***π*** and rate matrix **Q**. Scanning along an alignment from 5^’^ to 3^’^, the cumulative difference between any two clonal lineages accumulates as a sum of independent, identically distributed Bernoulli substitution trials. By Donsker’s invariance principle, as alignment length *L* → ∞, the standardized cumulative deviation curve:

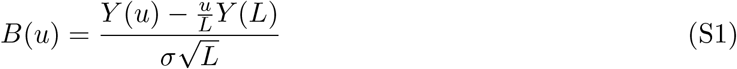

converges weakly to a standard Brownian bridge on *u* ∈ [0, 1], where E[*B*(*u*)] = 0 and Cov(*B*(*s*), *B*(*t*)) = min(*s, t*) − *st*. Under the null hypothesis of clonal evolution (*H*_0_), fluctuations in sequence affinity represent balanced thermal noise. Recombination is formally defined as an exogenous physical force that breaks this thermal equilibrium and induces a sustained, non-zero drift velocity 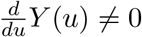 exceeding Brownian boundary-crossing thresholds.

### Assumption 3 (The Unpolarized Root Token as an Autapomorphic Sink)

Mutational rate heterogeneity (heterotachy) and directional hypermutation bursts (such as APOBEC, ADAR, or error-prone replication) frequently generate localized clusters of mutations along a single lineage. Heuristic triplet methods mistake these mutational clusters for horizontal donor acquisition. Reserving coordinate index 0 in the cross-attention simplex for an unpolarized ancestral root token **x**_root_ anchored at the metric origin **m**_root_ = **0** eliminates this vulnerability. In the cross-attention softmax:

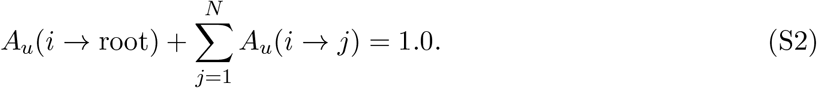

When lineage *i* acquires private substitutions that do not point toward any sampled contemporary taxon, the similarity logits to all contemporary leaves decrease uniformly and route attention into the root sink at index 0. Consequently, autapomorphic bursts reduce contemporary affinity without surging candidate donor attention. Routing private noise into the root sink provides intrinsic mathematical immunity to false positive alarms under rate heterogeneity.

### Assumption 4 (Spectral Landmark Representation and Iterative Tract Scrubbing on Simplex Δ^*K*^)

To scale linearly across large genomic cohorts (*N >* 1,000 genomes), the evolutionary variance of a population is represented by *K* spectral landmark centroids (*K* ≪*N* ) on the probability simplex Δ^*K*^. Landmarks are initialized via Farthest-Point Sampling (Gonzalez, 1985) on global distance matrix **D**_glob_ to ensure uniform coverage across deep clades, and are progressively purified through Iterative Forward Scrubbing (gap-masking detected recombinant tracts [*u*_1_, *u*_2_] in descending order of excursion amplitude so that masked positions exert zero force, Δ*F* = 0) followed by Backward Leave-One-In Verification. Every contemporary genome *i* is tracked as a continuous barycentric coordinate vector ***α***_*i*_(*u*) ∈ Δ^*K*^. Landmark projection and tract scrubbing reduce computational complexity from *O*(*N* ^2^*L*) or *O*(*N* ^3^*L*) down to *O*(*NKL*) while eliminating multi-event interference and parent-child polarity inversions.

### Physical Limits and Resolution Boundaries

The resolution of trajectory deconvolution is governed by fundamental physical boundaries:

1. *Mutational Density Floor* : Recombination can only be inferred if the introgressed tract contains distinguishing polymorphic mutations between parental lineages (*n*_mut_ = *L*_tract_ · *d*_parent_ ≥ 1). In low-divergence regimes (*d*_parent_ *<* 0.005), statistical power is fundamentally bounded by mutation availability rather than algorithmic sensitivity.
2. *Tract Length Resolution*: The dyadic scale-space matched filter operates across dyadic scales *τ* ∈ {15, 25, 50, 100, 200, 500} nt. Replacement tracts shorter than 15 nt (single-codon swaps) cannot be reliably distinguished from multi-nucleotide mutations without prior structural constraints.
3. *Extinction / Ghost Limits*: When an introgressing donor has no sampled relatives within phylogenetic distance *d*_max_, the tract is classified as an orphan ghost introgression. Directional consistency constraints prevent false attribution to contemporary outgroups, but donor clade identification remains impossible without surrogate sampling.

## S2 Complete Algorithmic Pipeline and Mathematical Formulations

### S2.1 Lossless SNP Compression and Prefix Distance Tensor

Given a multiple sequence alignment **M** ∈ Σ^*N*×*L*^ over alphabet Σ = {*A, C, G, T*, −}, columns exhibiting zero polymorphic variation across all *N* taxa carry zero phylogenetic information. Indexing out all monomorphic positions compresses alignment **M** into a segregating variant matrix 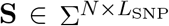 with bijective coordinate mapping function *Ψ* : {0, …, *L*_SNP_ − 1} → {0, …, *L* − 1}. For whole-genome bacterial alignments where *L*_SNP_ ≪ *L* (1,465-fold compression in *S. pneumoniae*), SNP indexing allows all trajectory computations to execute in memory-resident cache.

Across the compressed alignment, we compute the cumulative prefix difference tensor 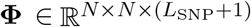.

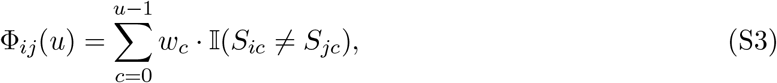

where *w*_*c*_ denotes site-specific weights (e.g., transition/transversion corrections or codon degeneracy). For any genomic sub-interval [*u*_1_, *u*_2_), the exact pairwise evolutionary distance between sequence *i* and sequence *j* is computed in *O*(1) time:

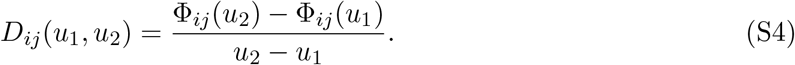

### S2.2 Global Divergence Matrix and Classical MDS Basis

The whole-sequence evolutionary divergence matrix **D**_glob_ ∈ ℝ^*N* ×*N*^ is evaluated across the full locus: *D*_glob,*ij*_ = Φ_*ij*_(*L*_SNP_)*/L*_SNP_. We construct the centering projection matrix 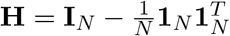 and form the Gower double-centered inner product matrix:

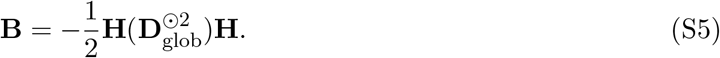

Spectral decomposition yields **B** = **VΛV**^*T*^, where eigenvalues are sorted in descending order *λ*_1_ ≥ · · · ≥ *λ*_*N*_ ≥ 0. We retain the top *k* = min(3, *N* − 1) positive eigenvalues **Λ**_*k*_ = diag(*λ*_1_, …, *λ*_*k*_) and corresponding orthonormal eigenvectors **V**_*k*_ ∈ ℝ^*N* ×*k*^ . The canonical basis is computed once and held fixed for the entire alignment to guarantee zero Procrustes rotational drift.

### S2.3 Exact Gower Out-of-Sample Continuous Trajectory Projection

At any genomic coordinate *u* ∈ [0, *L*_SNP_), each sequence *i* possesses an instantaneous local distance vector **d**_*i*_(*u*) ∈ ℝ^*N*^ to all reference taxa, evaluated over a symmetric Gaussian kernel window:

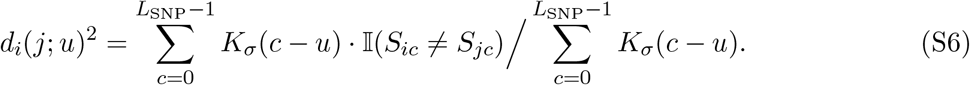

The exact Gower out-of-sample projection maps this local distance vector into canonical Euclidean space ℝ^*k*^ :

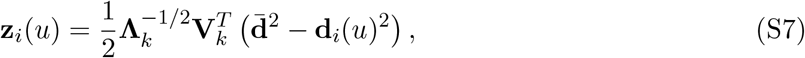

where 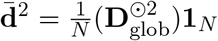 is the precomputed vector of average squared global distances. Evaluating **z**_*i*_(*u*) across all *u* produces the continuous trajectory tensor 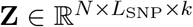 .

### S2.4 Continuous Phylogenetic Cross-Attention with Tree-RoPE and PhyloBias

To track affinity dynamics against reference landmarks without heuristic window slicing, `RhizAeon` employs continuous cross-attention. For each position *u*, sequence tokens are embedded into query (**q**_*i*_(*u*)), key (**k**_*j*_(*u*)), and value (**v**_*j*_(*u*)) representations in ℝ^*d*^. Phylogenetic distance constraints are injected through two mechanisms:

1. *Tree-RoPE (Tree Rotary Position Embedding)*: Metric MDS coordinates **m**_*i*_ ∈ ℝ^4^ modulate query and key vectors via multidimensional Givens rotations:

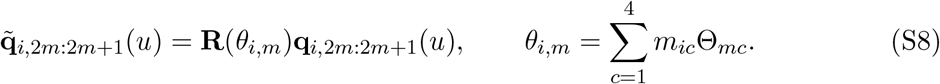
2. *PhyloBias*: A log-space continuous-time Markov substitution prior penalizes attention between divergent lineages:

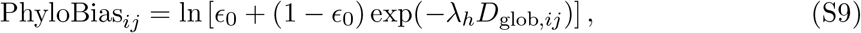

where *E*_0_ = 0.25 represents the equilibrium neutral substitution floor.

The resulting attention tensor **A** ∈ ℝ^*L*×*N* ×(*N* +1)^ evaluates attention from each query sequence *i* to all reference taxa *j* ∈ {1, …, *N* } and the root sink at index 0:

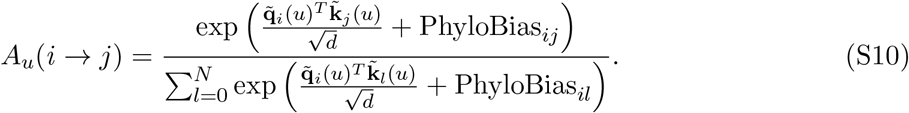

### S2.5 Cumulative Attention Curves and Differential Drift

For each sequence *i* and reference landmark bundle *m*, instantaneous attention is accumulated along the chromosome in *O*(*L*) time:

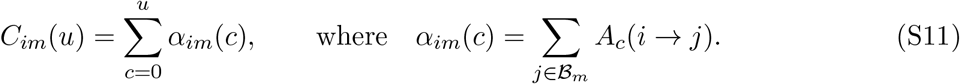

The home clade ℬ_home_(*i*) is identified as the majority recipient across the full locus: ℬ_home_(*i*) = arg max_*m*_ *C*_*im*_(*L*). The differential attention curve against candidate donor *m* is formulated as:

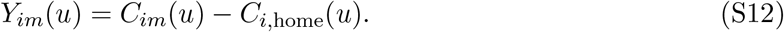

Under clonal evolution, sequence *i* maintains stable affinity to its home clade and exhibits negative or stationary drift *Y*_*im*_(*u*) ≤ 0. A recombination event from donor *m* produces a positive velocity excursion where 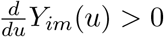.

### S2.6 Multi-Scale Dyadic Scale-Space Matched Filter and Running Drawdown

To detect both focal micro-conversions and macroscopic recombinant cassettes without parameter tuning, differential drift increments Δ*Y*_*im*_(*u*) = *Y*_*im*_(*u*) − *Y*_*im*_(*u* − 1) are convolved with zero-mean bipolar matched filter wavelets across dyadic scales *τ* ∈ {15, 25, 50, 100, 200, 500} nt:

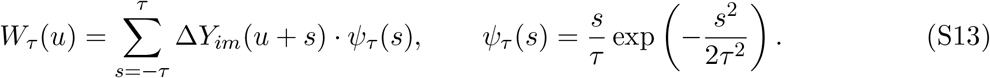

Concurrently, macroscopic cassettes are identified via Taylor running drawdown:

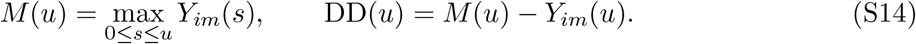

A candidate cassette is detected when the excursion drawdown exceeds the extreme-value threshold *Z*_phys_ ≥ *Z*_crit_, where 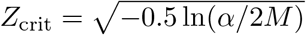 by Kolmogorov supremum boundary crossing.

### S2.7 Profile Maximum-Likelihood Breakpoint Polisher

Candidate changepoint intervals [*u*_1_ − *δ, u*_1_ + *δ*] and [*u*_2_ − *δ, u*_2_ + *δ*] localized by the matched filter are polished to single-nucleotide resolution using profile maximum likelihood. For each candidate coordinate *b*, the alignment is cleaved into left ([0, *b*)) and right ([*b, L*)) partitions. The profile likelihood ratio:

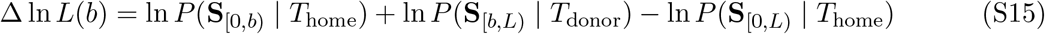

is evaluated under HKY85 substitution. The exact breakpoint 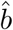 is localized at the maximum likelihood peak: 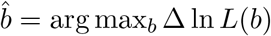. Significance is certified against the Dunn-Šidák corrected family-wise threshold *α*_FWER_ = 1 − (1 − *α*)^1*/M*^ .

## S3 Ancestral Recombination Graph (ARG) Reconstruction and Validation

### S3.1 Ancestral Event Consolidation

Raw recombinant tracts detected across contemporary genomes *ε*_raw_ = {*e*_1_, …, *e*_*M*_ } are clustered to distinguish ancestral gene transfers from recurrent hotspot activity. Two events *e*_*a*_ and *e*_*b*_ are grouped into ancestral cluster *C* if they share compatible donor clades and satisfy spatial boundary tolerance:

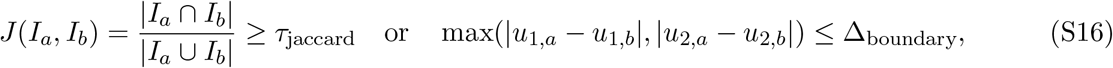

where default thresholds are *τ*_jaccard_ = 0.60 and Δ_boundary_ = 100 nt. The ancestral event coordinates are defined by median breakpoints *ū*_1_ = median({*u*_1,*i*_}) and *ū*_2_ = median({*u*_2,*i*_}). When |*C*| *>* 1, the transfer is parameterized as a shared ancestral reticulation node ancestral to all member isolates. When |*C*| = 1, it is parameterized as a private terminal event.

### S3.2 Structural Model Classes

A candidate evolutionary history *H* is parameterized by an ordered partitioning of the alignment [0, *L*) into *K* intervals 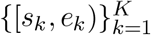 and an associated graph topology:

1. *Clonal Null* (*H*_0_): A single unpartitioned tree across the full locus (*K* = 1), requiring 2*N* − 3 branch lengths and 4 substitution parameters (*k* = 2*N* + 1).
2. *Unpolarized Partitioned Trees* (*H*_partition_): Independent unconstrained trees across all *K* partitions, requiring *k* = *K*(2*N* − 3) + 4*K* free parameters.
3. *Directed Polarized ARG* (*H*_ARG_): A single ancestral backbone tree where reticulations are represented as directed transfer edges that re-attach the recombinant clade inside the transfer interval [*u*_1_, *u*_2_]. Each reticulation edge adds only 2 free parameters (source node and target branch attachment) rather than 2*N* − 3 unconstrained branch lengths, which yields *k* = *K*_backbone_(2*N* − 3) + 2| *ε*_retic_| + 4.
4. *Modular Functional Architectures*: Partition models defined by structural protein domains (e.g., Spike NTD, ACE2 receptor-binding domain, and S2 fusion core).

### S3.3 Multidimensional Statistical Validation Engine

Competing evolutionary histories are evaluated across four orthogonal statistical criteria:

1. *Phylogenetic Profile Likelihood and Information Criteria*: Across each partition *k* ∈ {1, …, *K*}, log-likelihood ln *L*_*k*_ = ln *P* (**Y**_*k*_ | *T*_*k*_, **Q, *π***) is computed via vectorized Felsenstein pruning. Total log-likelihood ln *L* = Σ_*k*_ ln *L*_*k*_ is penalized under finite-sample corrected AICc and BIC:

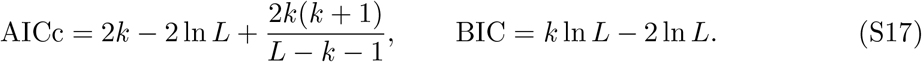
2. *Residual Non-Tree Manifold Strain*: The expected distance matrix is the length-weighted mixture of local partition patristic distance matrices:

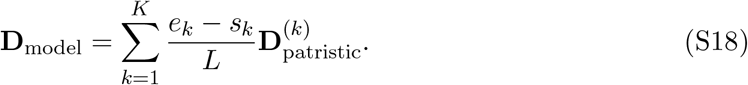

The residual non-tree distortion matrix **R** = **D**_obs_ − **D**_model_ yields the residual strain ratio:

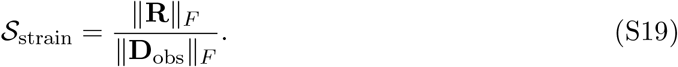

The percentage of non-tree strain explained by history *H* is (1 − *S*_strain_) × 100%.
3. *Spectral SNR over Random Matrix Bulk Edge*: Matrix **R** is centered via double-centering 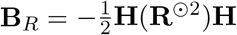. The leading eigenvalue *s*_1_ = *λ*_max_(**B**_*R*_) is compared against the theoretical Marchenko-Pastur bulk noise edge:

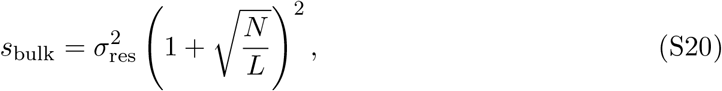

where *Σ*_res_ is the standard deviation of off-diagonal elements in **R**. An empirical SNR = *s*_1_*/s*_bulk_ ≤ 1.0 certifies that residual strain has collapsed into the bulk noise distribution.
4. *Synapomorphic Site Concordance Ratio (SSCR) and Polarity*. At each polymorphic site *u* where donor *D* and home *H* carry distinct nucleotides (*D*_*u*_ ≠ *H*_*u*_), recombinant allele *R*_*u*_ is scored as donor-concordant (*R*_*u*_ = *D*_*u*_) or home-concordant (*R*_*u*_ = *H*_*u*_):

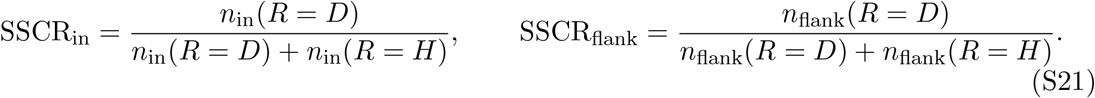

Polarization contrast is ΔSSCR = SSCR_in_ − SSCR_flank_. Statistical significance is evaluated via a one-sided Fisher’s exact test on [[*n*_in_(*D*), *n*_in_(*H*)], [*n*_flank_(*D*), *n*_flank_(*H*)]].

## S4 Coalescent ARG Simulation Protocol

Genealogies and ancestral recombination graphs (ARGs) were simulated using Hudson’s continuous-time coalescent framework with homologous recombination (Hudson, 1983; Posada and Crandall, 2001). For a sample of *n* = 10 contemporary lineages across an alignment of length *L* = 1,000 nucleotides, the coalescent process tracks active ancestral segments backward in time from *t* = 0. At any time *t*, let *k* denote the number of active ancestral chromosomes, and let *g*_*i*_ denote the genomic span between the leftmost and rightmost ancestral segments carried by lineage *i*, with total ancestral span 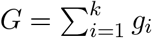.

The waiting time Δ*t* to the next evolutionary event is exponentially distributed with total event rate *λ*_tot_ = *λ*_coal_ + *λ*_rec_, where:

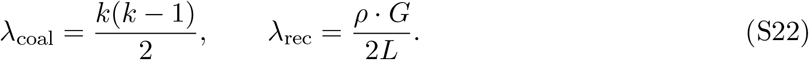

The population recombination parameter is defined as *ρ* = 4*N*_*e*_*r*, where *N*_*e*_ is the effective diploid population size and *r* is the per-generation recombination probability across the *L* nucleotide locus. With probability *λ*_coal_*/λ*_tot_, a coalescence event merges two randomly chosen lineages into an ancestral node. With probability *λ*_rec_*/λ*_tot_, a recombination event selects an eligible lineage with probability proportional to its span *g*_*i*_ and cleaves it at a uniform breakpoint coordinate *b* ∈ [*u*_start_, *u*_end_] to partition ancestral material into left and right lineages. Coalescent tracking halts when all *L* nucleotide sites reach their respective MRCA and yields an ordered sequence of *P* genealogical partitions, each defined by an interval [*s*_*p*_, *e*_*p*_) and a local gene tree *T*_*p*_.

Nucleotide sequences were evolved along the branch lengths of each local gene tree *T*_*p*_ under the continuous-time Markov HKY85 substitution model (Hasegawa et al., 1985). The model parameterizes unequal base frequencies ***π*** = [0.40, 0.20, 0.30, 0.10] and transition/transversion bias *κ* = 2.0. The infinitesimal generator matrix **Q** = (*Q*_*ij*_) is defined by:

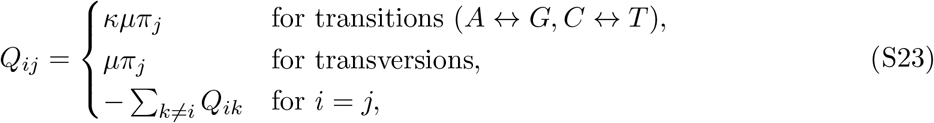

normalized such that average substitution rate − _*i*_ *π*_*i*_*Q*_*ii*_ = 1.0. Branch lengths *t*_*b*_ generated in coalescent units were scaled by the population mutation parameter *θ* = 4*N*_*e*_*µ* ∈ {10, 50, 100, 200} so that the effective branch length is *b* = *t*_*b*_ · *θ/*(2*L*). Site-rate heterogeneity was introduced by drawing independent site mutation rate multipliers *r*_*u*_ ∼ Gamma(*α*, 1*/α*) across *u* ∈ [0, *L*) such that E[*r*_*u*_] = 1.0 and Var(*r*_*u*_) = 1*/α* across shape parameters *α* ∈ {2.0, 0.5, 0.05}.

## S5 Specialized Epidemiological and Genomic Simulation Suites

Four specialized simulation generators probed operational regimes outside the neutral coalescent (Figure 2):

- *Epidemic Clonal Expansion*: Trees were constructed by generating a bifurcating clonal backbone of 40 background parent taxa across three parental divergence levels (*d*_parent_ ∈ {0.02, 0.05, 0.10}). An ancestral recombination event transferred a 1,501 nt cassette (coordinates 1,500–3,000) from Parent 2 to an ancestral recipient in Parent 1 to form recombinant ancestor *R*_anc_. From *R*_anc_, a burst of *K* ∈ {2, 5, 8, 10, 15} descendant lineages was generated with short terminal branch divergence (*d*_swarm_ = 0.001) to model transmission swarms in clinical cohorts (*N* ∈ {42, 45, 48, 50, 55}, 20 replicates each, 300 alignments, *L* = 5,000 nt, *K*_true_ = 1).
- *Multi-Way Mosaic Swarms*: Backbones comprised 45 background parent taxa evenly divided among three distinct parental clades (15 taxa each for P1, P2, P3) at *d*_parent_ ∈ {0.02, 0.05, 0.10}. Three multi-way topologies were simulated. In the *Dual Cassette* regime, recombinant lineages acquired a 1,200 nt cassette from P2 (coordinates 1,000–2,200) and a separate 1,200 nt cassette from P3 (coordinates 3,400–4,600) embedded within a P1 background (*L* = 5,000 nt). In the *Nested Russian Doll* regime, an outer 3,000 nt cassette from P2 (coordinates 1,000– 4,000) embedded an inner 1,000 nt cassette from P3 (coordinates 2,000–3,000) to produce a hierarchical *P*_1_ → *P*_2_ → *P*_3_ → *P*_2_ → *P*_1_ mosaic (*L* = 6,000 nt). In the *Triple Crossover Cascade* regime, sequential crossovers produced a serial chromosome mosaic composed of P1 (1–2,000 nt), P2 (2,001–4,000 nt), and P3 (4,001–6,000 nt) across *L* = 6,000 nt. Descendant swarms spanned *K* ∈ {1, 5, 10} (20 replicates each, 540 alignments).
- *Ghost Donor Introgression*: Alignments of length *L* = 5,000 nt contained 30–40 background parent taxa across three ghost donor regimes (*d*_parent_ ∈ {0.02, 0.05, 0.10}, *K* ∈ {2, 5, 10}, 20 replicates each, 540 alignments). In the *Surrogate Sister Clade* regime, an ancestral lineage diverged into donor *D*_ghost_ and surrogate sister 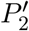, after which *D*_ghost_ transferred cassette [1,500, 3,000] to *P*_1_ before omission while sister 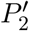 remained in the alignment. In the *Pure Ghost Lineage* regime, the donor lineage branched from the tree root and transferred cassette [1,500, 3,000] into *P*_1_ before complete omission so that only unrelated contemporary taxa remained in the sample. In the *Deep Zoonotic Reservoir* regime, the introgressing tract originated from an outgroup lineage with cross-species evolutionary distance *d*_reservoir_ = 0.15 into *P*_1_.
- *Bacterial Micro-Conversions*: Alignments (*L* = 5,000 nt) contained 30 background parent taxa and recombinant swarms (*K* ∈ {2, 5, 10}) across five conversion tract lengths (*L*_tract_ ∈ {50, 100, 200, 400, 800} nt, inserted at coordinate 2,100) and three divergence levels (*d*_parent_ ∈ {0.02, 0.05, 0.10}, 20 replicates each, 900 alignments). The expected number of informative nucleotide substitutions per micro-tract ranged from 1–5 mutations at 50 nt to 16–74 mutations at 800 nt.

## S6 Grand-Scale Continuous Parameter Sweep (1,000,000 Genomes)

To evaluate algorithmic performance, false positive rate calibration, and multi-event mosaic resolution across unconstrained continuous evolutionary space rather than isolated grid points, we executed a grand continuous parameter sweep of 1,000,000 independent whole-genome alignments. The sweep comprised 333,334 non-recombinant clonal nulls and 666,666 recombinant chimeras (Table S3, Figures 3 and S1). Each trial sampled an independent parameter vector uniformly or log-uniformly across evolutionary phase space:

- *Alignment Length (L)*: Sampled log-uniformly from *L* ∼ LogUniform(300, 30,000) nt and discretized to multiples of 50 nt (*L* ∈ [300, 30,000] nt, median 5,450 nt).
- *Sample Taxon Count (N)*: Stratified across three cohort surveillance regimes: 40% small outbreak cohorts (*N* ∼ Uniform(4, 30) taxa), 40% medium surveillance cohorts (*N* ∼ Uniform(31, 100) taxa), and 20% large population cohorts (*N* ∼ Uniform(101, 250) taxa).
- *Parental Divergence (d*_parent_): Log-uniformly distributed across *d*_parent_ ∼ LogUniform(0.005, 0.350) substitutions per site.
- *Substitution Dynamics* : Continuous-time Markov substitution under the HKY85 model with transition/transversion ratio *κ* = 2.0 and empirical base frequencies ***π*** = [0.30, 0.20, 0.20, 0.30]. Position-specific finite-time transition matrices **P**(*t*) were precomputed analytically per trial for rapid, exact simulation without transcendental overhead.
- *Site Rate Heterogeneity and Biological Heterotachy (α)*:
  - *Strict Molecular Clock* (25% of trials, *N* = 250,000 overall, 83,311 in null partition): *α* = 1,000.0 (uniform substitution rates across all sites).
  - *Continuous Gamma Heterotachy* (75% of trials, *N* = 750,000 overall, 250,023 in null partition): Shape parameter drawn log-uniformly from *α* ∼ LogUniform(0.20, 5.00), with position-specific relative mutation rates *r*_*u*_ ∼ Gamma(*α, α*) sampled independently per site and clamped to [0.01, 50.0].
- *Lineage Architecture and Transmission Drift* : The ancestral root sequence evolved into parental nodes *P*_1_, *P*_2_, and optional *P*_3_ (for cohorts with *N* ≥ 8 taxa). Contemporary leaves accumulated post-recombination private mutations along terminal branches with transmission drift *d*_drift_ ∼ Uniform(0.0005, 0.0035) subs/site.
  - *Reticulation Cardinality (K*_true_):
  - *K*_true_ = 0 (33.33% of trials, *N* = 333,334): Strictly non-recombinant clonal nulls to calibrate empirical false positive rates under severe rate variation.
  - *K*_true_ = 1 (36.64% of trials, *N* = 366,396, 55.0% of recombinant partition): Single reticulation event per genome.
  - *K*_true_ = 2 (16.70% of trials, *N* = 166,974, 25.0% of recombinants): Dual independent recombination events across segmented partitions.
  - *K*_true_ = 3 (7.99% of trials, *N* = 79,881, 12.0% of recombinants): Triple independent recombination events (multi-way mosaicism).
  - *K*_true_ = 4 (5.34% of trials, *N* = 53,415, 8.0% of recombinants): Quadruple reticulations that include complex nested architectures.
- *Structural Event Topologies*:
  - *5’ Terminal Crossovers* (20% of terminal events): Breakpoint initiates at genome start (*u*_1_ = 1) and terminates at *u*_2_ ∼ Uniform(50, segment bound).
  - *3’ Terminal Crossovers* (20% of terminal events): Breakpoint initiates at *u*_1_ ∼ Uniform(segment start, *L*− 50) and extends to chromosome end (*u*_2_ = *L*).
  - *Intercalary Conversion Cassettes* (60% of events): Internal tracts (1 *< u*_1_ *< u*_2_ *< L*) with tract length *L*_tract_ sampled log-uniformly from 30 bp to 0.65 × segment length.
- *Multi-Donor Mosaicism (P*_1_ ← *P*_2_ + *P*_3_): In cohorts with *N* ≥ 8 taxa (*N* = 179,233 trials), recombinant chimeras were generated with multiple distinct donor clades where distinct segments of the same genome were inherited from independent ancestral lineages *P*_2_ and *P*_3_ relative to recipient background *P*_1_.

Simulations were implemented in native compiled Rust (`crates/rhizaeon-cli/src/bin/grand_sweep.rs`) leveraging Rayon data parallelism. The 1,000,000 trials were distributed across cluster infrastructure as a 50-task Slurm array (50 tasks × 20,000 trials, 32 CPU threads and 48 GB RAM per task across 512 distributed CPU cores). The benchmark maintained a sustained throughput of ≈ 380 complete whole-genome simulations and scans per second and completed all 1,000,000 trials in 43.8 minutes. Outputs were aggregated, deduplicated, and verified using `scripts/simulations/merge_sweep_1m.py`to ensure exact, reproducible execution across the entire corpus.

## S7 Ground Truth Extraction and Bijective Bipartite Matching

To establish ground truth for reticulation events without relying on heuristic approximations, we computed exact local patristic distance matrices **D**^(*p*)^ ∈ ℝ^*n*×*n*^ across every genealogical partition *p* ∈ {1, …, *P* } from the branch lengths of tree *T*_*p*_. For each contemporary taxon *i*, its local phylogenetic sister at nucleotide *u* ∈ [*s*_*p*_, *e*_*p*_) is the closest non-identical taxon:

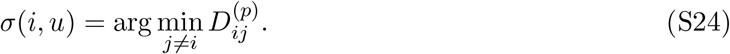

The background Home parent *H*_*i*_ is defined as the majority sister across the full sequence: *H*_*i*_ = mode_*u*∈[0,*L*)_*Σ*(*i, u*). A true reticulation event occurs when taxon *i*’s sister shifts to a distinct donor lineage *D* = *Σ*(*i, u*) ≠ *H*_*i*_ across a contiguous segment [*u*_1_, *u*_2_]. To eliminate trivial non-divergent ancestral coalescences, ground truth events were filtered to require at least one distinguishing mutation between Home and Donor across the tract 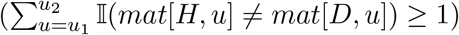.

To score inferred events against ground truth without bias, predicted events 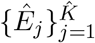 were paired to true events 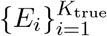 via maximum-weight bijective bipartite matching using the Hungarian algorithm (`scipy.optimize.linear_sum_assignment`). The pairwise affinity matrix *A*_*ij*_ ∈ [0, 1] between true event *i* = (*C*_*i*_, *H*_*i*_, *D*_*i*_, *u*_1,*i*_, *u*_2,*i*_) and inferred event 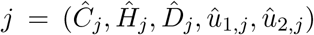 was formulated as:

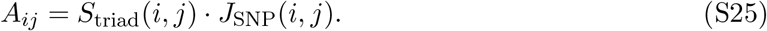

Here 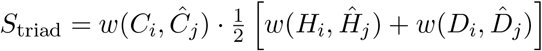 weights lineage attribution using tree patristic distances with Gaussian kernel width 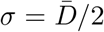. Spatial overlap *J*_SNP_ = |*S*_true_ ∩ *S*_inferred_|*/*|*S*_true_ ∪ *S*_inferred_| is the Jaccard index evaluated over distinguishing polymorphic sites between Home and Donor. Unmatched inferred events were penalized as false alarms (reducing precision PPV = 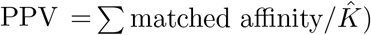), whereas unmatched true events reduced sensitivity (TPR = Σmatched affinity*/K*_true_).

## S8 Empirical Genomic Cohorts and Annotations

### Sarbecovirus Whole-Genome Cohort and Coordinate Frame Conversion

We analyzed two nested whole-genome alignments compiled from GISAID and GenBank. The primary 78-taxon sarbecovirus alignment of Lytras et al. (2022) spans 30,956 bp in multiple sequence alignment (MSA) coordinates with 13,550 total polymorphic columns and 13,160 gap-filtered segregating sites. An expanded 83-taxon alignment (30,965 bp MSA, 13,260 gap-filtered sites) incorporates the five Laotian *Rhinolophus* bat sarbecoviruses (BANAL-20-52, BANAL-20-103, BANAL-20-116, BANAL-20-236, and BANAL-20-247) reported by Temmam et al. (2022). The five Laotian BANAL genomes introduce a 9 nt insertion in the Spike N-terminal domain around 78-MSA position 22,050, which shifts all downstream alignment coordinates by +9 nt relative to the 78-taxon MSA. In their primary study, Lytras et al. (2022) reported open reading frame annotations and the 21 consensus GARD breakpoints (derived from a 22-virus subset) in both ungapped SARS-CoV-2 Wuhan-Hu-1 reference coordinates (NC 045512.2, 29,903 nt, columns `SC2_start`/`SC2_end` in their Supplementary Table S3) and 78-taxon MSA coordinates (columns `al start`/`al end`). To prevent cross-frame coordinate discrepancies, Table S1 provides the exact bijective conversion across all three coordinate systems: Wuhan-Hu-1 reference (1–29,903 nt), 78-taxon MSA (1–30,956 nt), and 83-taxon MSA (1–30,965 nt). The table maps all sarbecovirus open reading frames, functional domains, and recombination hotspots and coldspots. It also pairs all 21 published GARD breakpoints with the closest `RhizAeon` continuous changepoints evaluated within each matching frame.

### *Streptococcus pneumoniae* PMEN1 Whole Chromosome

We analyzed the complete 2.22 Mb chromosome of the pandemic multidrug-resistant Spain23F-1 / PMEN1 lineage of *S. pneumoniae* (ATCC 700669, 11 representative global isolates spanning 1984–2008, Croucher et al. 2011). Lossless SNP indexing identified 1,516 segregating polymorphic sites across the 2,221,315 bp chromosome. Annotations follow ATCC 700669 (FM211187.1) across resistance and virulence loci: *pbp2x* (nt 287,466–289,718), capsular polysaccharide locus *cps* (nt 334,103–354,291), *pbp1a* (nt 322,540– 324,702), *pbp2b* (nt 1,532,410–1,534,452), dihydrofolate reductase *folA* (nt 1,623,115–1,623,600), and surface antigen adhesin *pspC/cbpA* (nt 2,109,335–2,111,878).

### Hominoidea MHC Class I Allelic Cohort

We analyzed *N* = 379 classical MHC Class I alleles (1,128 bp coding alignment across human, chimpanzee, gorilla, and orangutan, Maccari et al. 2017). Alleles were retrieved from the IPD-IMGT/HLA and IPD-MHC databases, comprising 150 *HLA-A*, 150 *HLA-B*, and 50 *HLA-C* human alleles alongside 15 chimpanzee (*Patr* ), 8 gorilla (*Gogo*), and 6 orangutan (*Popy* ) alleles. Exon annotations follow canonical classical HLA Class I structure: Exon 1 (Leader peptide, nt 1–72), Exon 2 (*α*_1_ domain, nt 73–342), Exon 3 (*α*_2_ domain, nt 343–618), Exon 4 (*α*_3_ domain, nt 619–894), Exon 5 (Transmembrane anchor, nt 895–1,011), and Exons 6–7 (Cytoplasmic tail, nt 1,012–1,128).

**Table S1:**
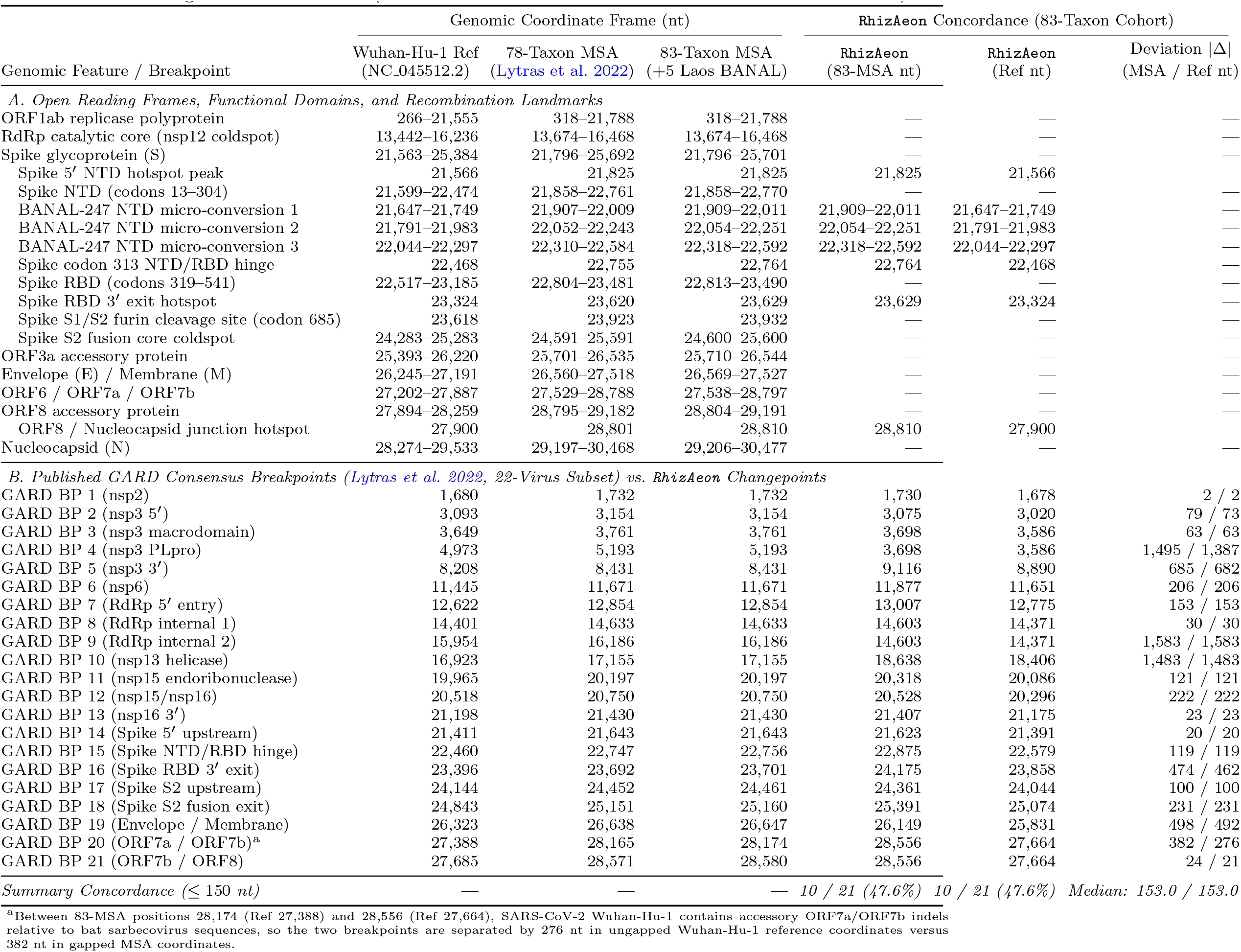
Bijective coordinate conversion across SARS-CoV-2 Wuhan-Hu-1 reference coordinates (NC 045512.2, 1–29,903 nt), the 78-taxon multiple sequence alignment of Lytras et al. (2022) (1–30,956 nt), and the expanded 83-taxon multiple sequence alignment incorporating the five Laotian BANAL isolates of Temmam et al. (2022) (1–30,965 nt). Part A lists major open reading frames, Spike functional domains, coldspots, hotspots, and BANAL-247 micro-conversion tracts. Part B lists all 21 consensus GARD breakpoints from Lytras et al. (2022) (Supplementary Table S3) alongside the closest `RhizAeon` changepoints and spatial deviations (|Δ|, nt) evaluated strictly within the matching coordinate frame (83-taxon MSA frame vs. Wuhan-Hu-1 reference frame).

**Table S2:**
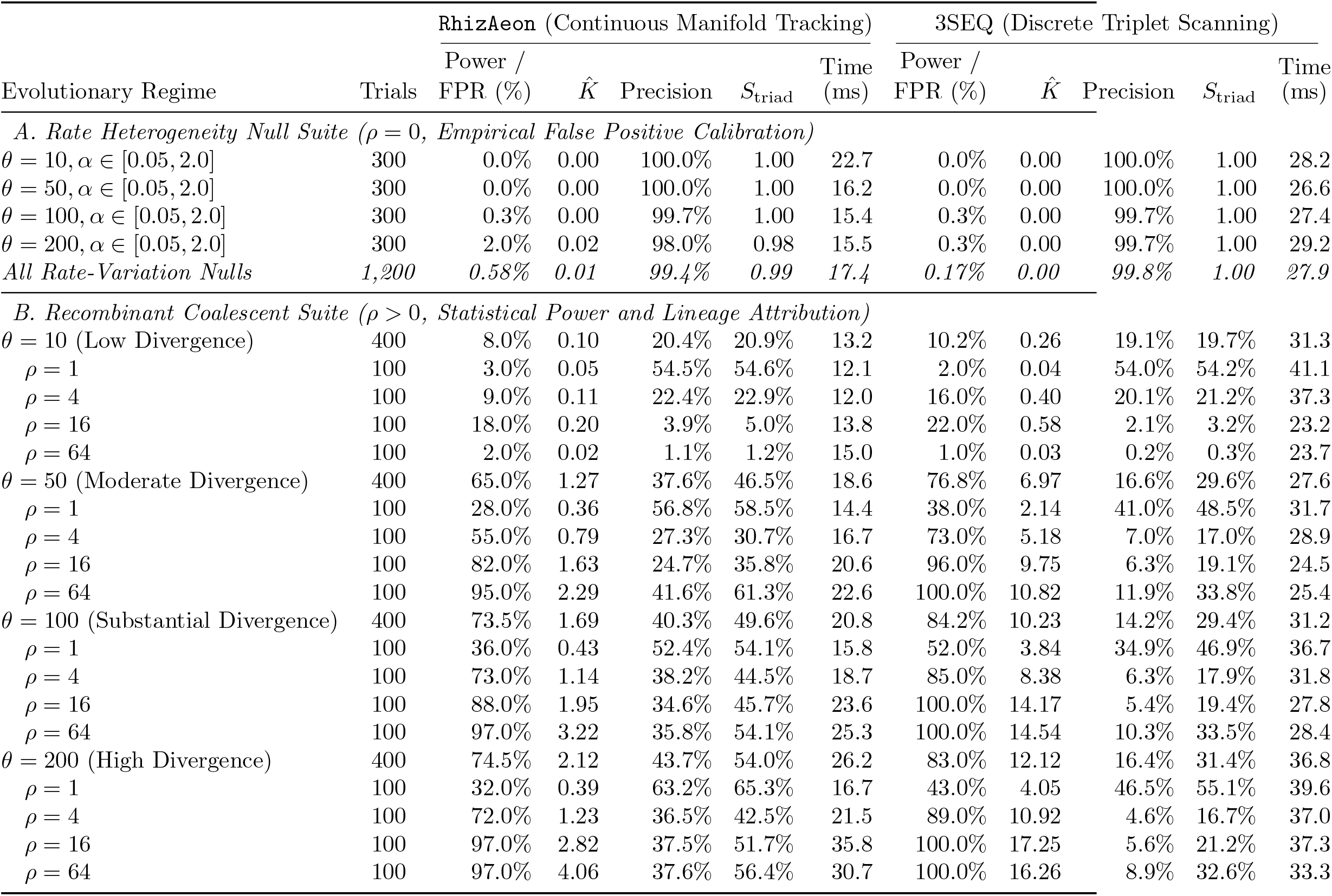
Quantitative comparison of `RhizAeon` and 3SEQ across the 3,200-trial Posada & Crandall (2001) benchmark suite. Results are grouped by evolutionary regime: false positive evaluation under Gamma rate heterogeneity (*α* ∈ {2.0, 0.5, 0.05}, *ρ* = 0, 1,200 trials), clock-like null controls (*ρ* = 0, 400 trials), and recombinant coalescent trials (*ρ* ∈ {1, 4, 16, 64}, 1,600 trials). Columns report the population mutation parameter (*θ*), population recombination parameter (*ρ*), binary alignment-level power or FPR (%), mean inferred event count 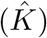, continuous bipartite event precision (PPV), triad lineage attribution accuracy (*S*_triad_), and execution runtime per alignment (ms). Whereas alignment-level power scores binary locus detection, PPV weights one-to-one Hungarian-matched events by *S*_triad_ × *J*_SNP_ and penalizes ungrounded or over-split calls (Section 3.1).

**Table S3:**
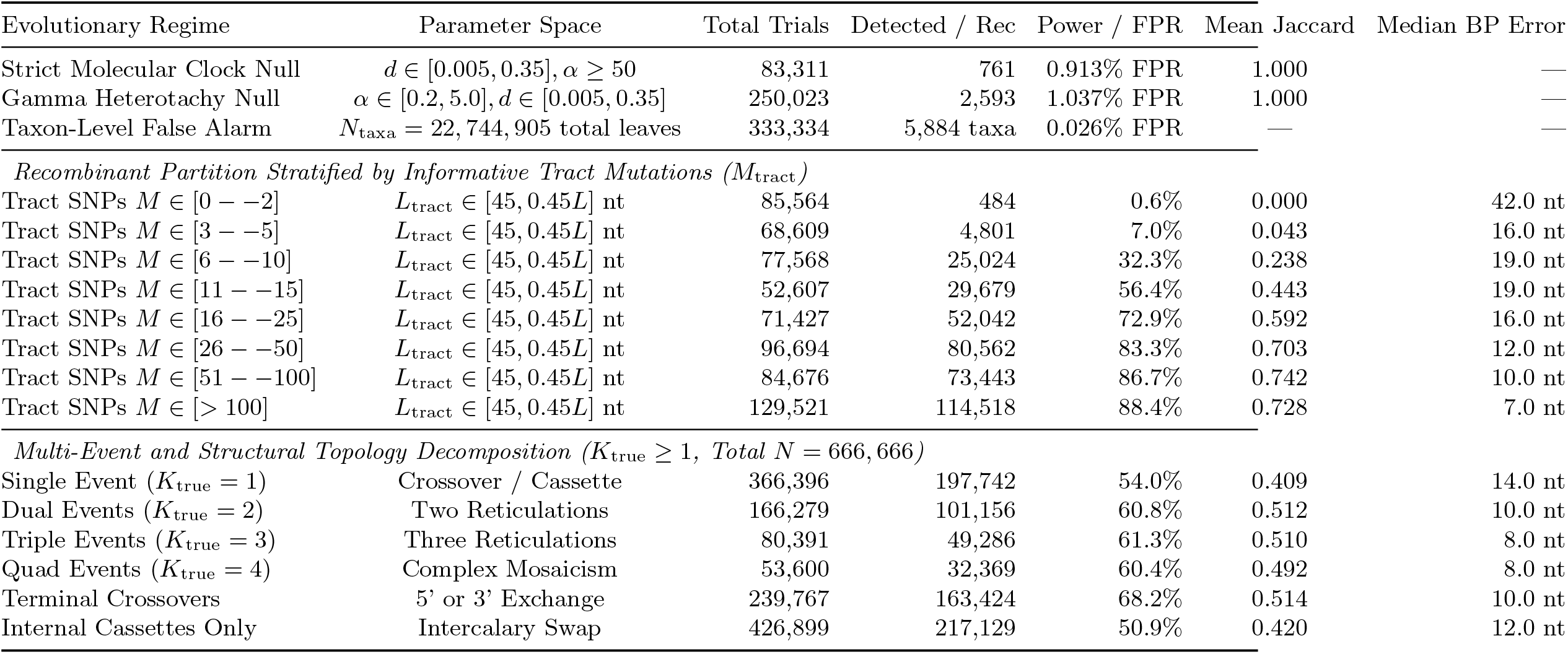
Empirical operational phase envelope of `RhizAeon` across 1,000,000 independent continuous simulation trials. Simulations span *L* ∈ [300, 30,000] nt, *N* ∈ [4, 250] taxa, *d*_parent_ ∈ [0.005, 0.35] subs/site, and continuous Gamma rate variation *α* ∈ [0.2, 5.0] alongside strict molecular clock controls (*α* ≥ 50). Multi-event topologies include true nulls (*K* = 0), 5’/3’ terminal crossovers, intercalary conversion cassettes (*K* = 1), and multi-donor / multi-event mosaicism (*K* ∈ {2, 3, 4}). Breakpoint errors report median absolute error (nt) among detected recombinant tracts.

**Figure S1:**
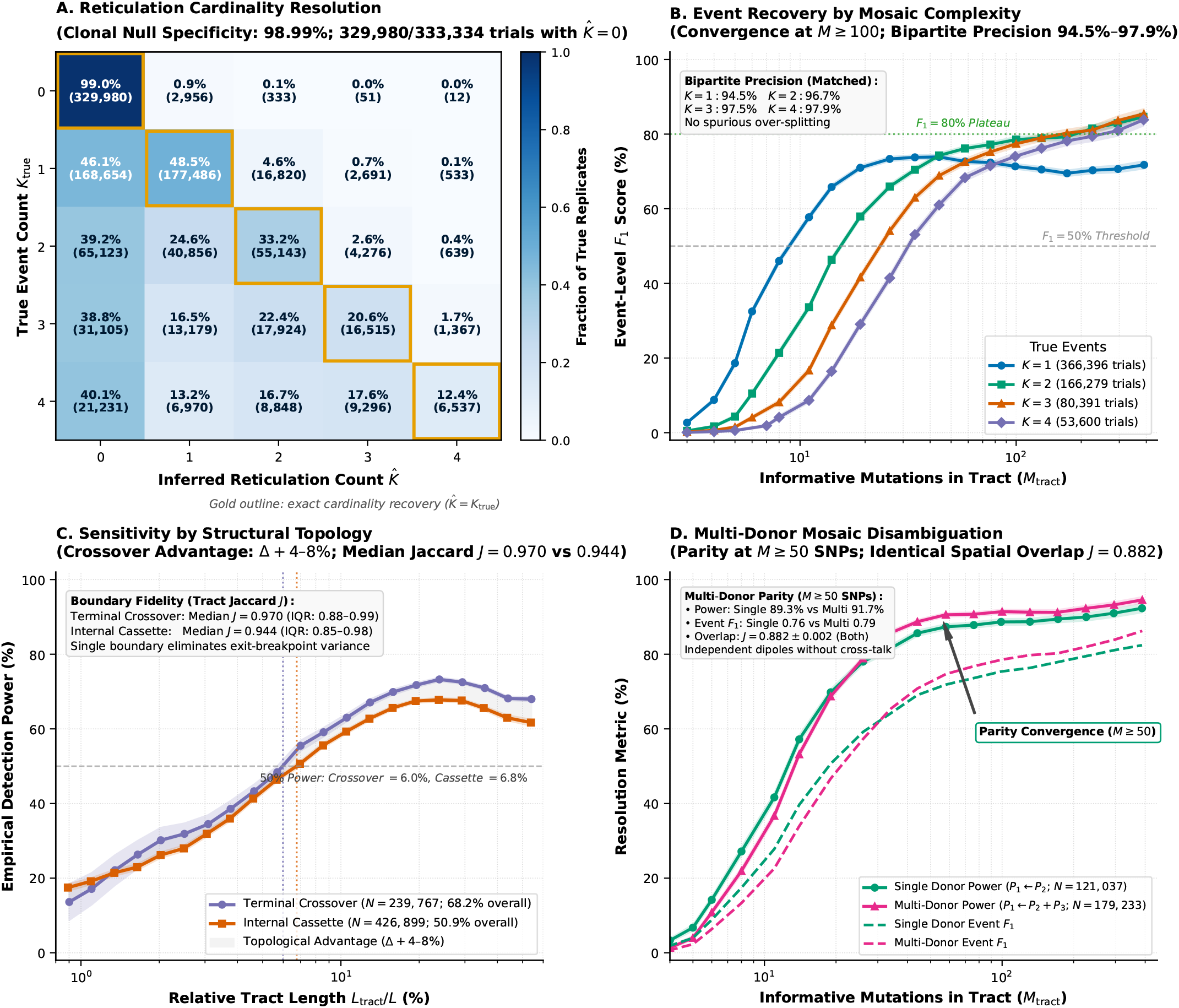
Multi-event reticulation resolution, structural topology sensitivity, and multi-donor disambiguation across the 1,000,000-trial simulation sweep. (A) Cardinality confusion matrix comparing true (*K*_true_) versus inferred 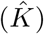 event counts across 333,334 clonal nulls (*K*_true_ = 0, 98.99% specificity, 1.01% alignment-level FPR) and 666,666 recombinant cohorts (*K*_true_ ∈ {1, 2, 3, 4}). (B) Event-level *F*_1_ score across informative tract mutations (*M*_tract_) stratified by *K*_true_ ∈ {1, 2, 3, 4} with 95% bootstrap confidence ribbons. (C) Detection power across relative tract length (*L*_tract_*/L*) comparing terminal crossovers (*N* = 239,767) against intercalary cassettes (*N* = 426,899). (D) Sensitivity and event *F*_1_ score across *M*_tract_ comparing single-donor (*P*_1_ ← *P*_2_) versus multi-donor (*P*_1_ ← *P*_2_ + *P*_3_) mosaics.

